# A Non-negative Measure Of Feature-Related Information Transfer Between Neural Signals

**DOI:** 10.1101/758128

**Authors:** Jan Bím, Vito De Feo, Daniel Chicharro, Malte Bieler, Ileana L. Hanganu-Opatz, Andrea Brovelli, Stefano Panzeri

**Author notes:** Corresponding authors: Stefano Panzeri Andrea Brovelli. Co-senior authors (Andrea Brovelli, Stefano Panzeri).

## Abstract

Quantifying both the amount and content of the information transferred between neuronal populations is crucial to understand brain functions. Traditional data-driven methods based on Wiener-Granger causality quantify information transferred between neuronal signals, but do not reveal whether transmission of information refers to one specific feature of external stimuli or another. Here, we developed a new measure called Feature-specific Information Transfer (FIT), that quantifies the amount of information transferred between neuronal signals about specific stimulus features. The FIT quantifies the feature-related information carried by a receiver that was previously carried by a sender, but that was never carried by the receiver earlier. We tested the FIT on simulated data in various scenarios. We found that, unlike previous measures, FIT successfully disambiguated genuine feature-specific communication from non-feature specific communication, from external confounding inputs and synergistic interactions. Moreover, the FIT had enhanced temporal sensitivity that facilitates the estimation of the directionality of transfer and the communication delay between neuronal signals. We validated the FIT’s ability to track feature-specific information flow using neurophysiological data. In human electroencephalographic data acquired during a face detection task, the FIT demonstrated that information about the eye in face pictures flowed from the hemisphere contralateral to the eye to the ipsilateral one. In multi-unit activity recorded from thalamic nuclei and primary sensory cortices of rats during multimodal stimulation, FIT, unlike Wiener-Granger methods, credibly detected both the direction of information flow and the sensory features about which information was transmitted. In human cortical high-gamma activity recorded with magnetoencephalography during visuomotor mapping, FIT showed that visuomotor-related information flowed from superior parietal to premotor areas. Our work suggests that the FIT measure has the potential to uncover previously hidden feature-specific information transfer in neuronal recordings and to provide a better understanding of brain communication.

**Author summary:** The emergence of coherent percepts and behavior relies on the processing and flow of information about sensory features, such as the color or shape of an object, across different areas of the brain. To understand how computations within the brain lead to the emergence of these functions, we need to map the flow of information about each specific feature. Traditional methods, such as those based on Wiener-Granger causality, quantify whether information is transmitted from one brain area to another, but do not reveal if the information being transmitted is about a certain feature or another feature. Here, we develop a new mathematical technique for the analysis of brain activity recordings, called Feature-specific Information Transfer (FIT), that can reveal not only if any information is being transmitted across areas, but whether or not such transmitted information is about a certain sensory feature. We validate the method with both simulated and real neuronal data, showing its power in detecting the presence of feature-specific information transmission, as well as the timing and directionality of this transfer. This work provides a tool of high potential significance to map sensory information processing in the brain.

## Introduction

Cognitive functions, such as the generation of conscious percepts or the selection of appropriate actions in changing sensory environments, emerge from the dynamic processing and routing of information about sensory features across different areas of the brain (Bressler and Menon, 2010; Runyan et al., 2017; Van Vugt et al., 2018; Varela et al., 2001). Tracking such feature-specific flow of information across both microscopic and macroscopic neuronal networks is therefore crucial for modern neuroscience.

To quantify the degree of communication between neural populations or brain areas from the statistical dependencies between neural signals (Bressler and Seth, 2011; Brovelli et al., 2004; Seth et al., 2015), the most successful model-free methods rely on the Wiener-Granger principle (Granger, 1969; Wiener, 1956). This principle identifies information flow between time series when future values of a given signal can be predicted from the past values of another, above and beyond what can be achieved from its autocorrelation. The information theoretic measures based on the Wiener-Granger principle, Transfer Entropy (Schreiber, 2000) and Directed Information (Massey, 1990), represent the most general measures of Wiener-Granger causality and capture any (linear and nonlinear) time-lagged conditional dependence between neural signals (Besserve et al., 2015; Vicente et al., 2011).

Such measures have been instrumental in untangling many aspects of brain communication (see e.g. (Besserve et al., 2015; Bosman et al., 2012; Brovelli et al., 2004; Li et al., 2019; Stramaglia et al., 2016; Van Kerkoerle et al., 2014; Vinck et al., 2015)). However, they do not cast light on the information content of brain communication. That is, these Wiener-Granger measures do not reveal what type of information is carried, that is whether the information transferred from one area to the next is about a certain sensory feature (for example, the color or shape of an object) or another feature.

Here, we exploit recent progress in Partial Information Decomposition (PID, see (Wibral et al., 2017; Williams and Beer, 2010)) to develop a novel non-negative measure of feature-specific information transfer that can reveal not only if any information is being transmitted across areas, but whether or not such information is about a certain sensory feature. The PID framework (Wibral et al., 2017; Williams and Beer, 2010) allows to decompose the information about a target variable that several sources provide individually (unique information), in a shared way (redundant information), or jointly by their combination (synergistic information). Most importantly, these terms are non-negative. Within this framework, we propose a novel metric named Feature-specific Information Transfer (FIT), which formalizes feature-related information transfer in a way that incorporates both the Wiener-Granger principle (Granger, 1969; Wiener, 1956) and the redundancy/synergy principles of neural population coding (Pola et al., 2003; Schneidman et al., 2003; Williams and Beer, 2010). In intuitive terms, our measure identifies the part of information about a stimulus feature encoded in the current activity of a receiver, which was not encoded in the past activity of the receiver, and which was instead encoded by the past activity of the sender. We investigate the relevance of the proposed metric on simulated data which illustrate different scenarios of information transfer between two signals. Then, we test FIT’s ability to track feature-specific information flow from neurophysiological data recorded from human participants in cognitive settings (EEG and MEG) and from rats (multi-unit activity) during multisensory stimulations.

## Results

### Feature-specific Information Transfer (FIT)

Consider two simultaneously-recorded neural signals *X* (a sender of information) and *Y* (a receiver of information), both of which may carry information about a feature *S*. This feature may be any experimental variable under investigation, such as the type of sensory stimulus being presented, the behavioral response or any other task variable. The considered neural signals may be, for example, the spiking activity of single or multiple neurons, or the aggregate activity of neural populations or areas recorded with neurophysiological and neuroimaging techniques, such as EEG, MEG or fMRI.

Established information theoretic methods relying on the Wiener-Granger principle, such as Directed Information (DI) (Massey, 1990), quantify the information transmitted from *X* to *Y* as the mutual information between the neural activity *Y*_*t*_ of the receiver at time t and the activity *X*_*t*−*d*_ of the sender in the past lagged by a time delay *d*, conditioned upon the past activity *Y*_*t*−*d*_ of the receiver (see Eq. 1 of Methods). The DI does not make any reference to the modulation of neural activity by a feature of the external stimulus, and thus it reflects all sources of information transfer, for example about either sensory features or internal state variables.

To quantify information that is transmitted specifically about a certain stimulus feature, we developed a new information theoretic measure that we termed the Feature-specific Information Transfer (FIT). Building on the PID framework (Williams and Beer, 2010), FIT is defined as information about the stimulus feature *S*, carried by the present activity of a receiver *Y*_*t*_ at time t, that is redundant with the information about *S* carried by the past activity *X*_*t*−*d*_ of the sender, and which at the same time is also unique, or novel, with respect to the information about *S* carried the past activity *Y*_*t*−*d*_ of the receiver. These requirements are intuitively important to identify genuine transmission of feature information from *X* to *Y*. If feature information in *Y* is genuinely received from *X*, then it must be that the feature information in *X* was present in the same form earlier (hence requirement of redundancy of feature information in *X* at a previous time) and that it has not been present in Y earlier (hence the requirement of uniqueness of present feature information in Y with respect to past information in Y). We refer to Materials and Methods for details about the mathematical derivation of the FIT.

One previous study has attempted to define a measure of feature-specific transfer, called Directed Feature Information (DFI), see (Ince et al., 2015). The DFI was also built on the requirement of redundancy of feature information in the past of X and novelty with respect to the past of Y. However, the DFI had two major differences with respect to the FIT: i) the DFI used a definition of redundancy which does not isolate only redundancy, but rather includes synergistic effects (and because of this the DFI can be negative, a property that poses questions regarding its suitability as a general measure of feature-specific information transfer, see (Ince et al., 2015)); and ii) the DFI discounts past information in X by simply conditioning upon its past values rather than removing all non-unique information.

In principle, FIT has two key advantages with respect to DI and DFI. Firstly, unlike for DI, FIT focuses on feature-specific information, neglecting feature-unspecific information. Secondly, the FIT is expected to discount more conservatively spurious effects of autocorrelations or previous information in the receiving signal than achieved by DI and DFI, because it is built on the concept of retaining only feature information that is unique with respect to that carried by the past activity of the receiver (discarding all non-unique information), rather than simply on conditioning on the past activity of the receiver (as done with the DI and DFI).

To test these expectations and assess the potentially powerful properties of the FIT measure in empirical analyses, we performed numerical simulations capturing three different scenarios of stimulus-specific information transfer that could occur between neural populations (Fig. 1). In all three scenarios in Fig. 1, information was transmitted from X to Y with 30 ms delay in the period from 40 to 60 ms. We simulated information transmission to be precisely localized in time, so to test the temporal precision of the various measures in detecting information transfer. What varied between the three scenarios was if and how feature-related information was transferred from *X* to *Y*.

**Figure 1.**
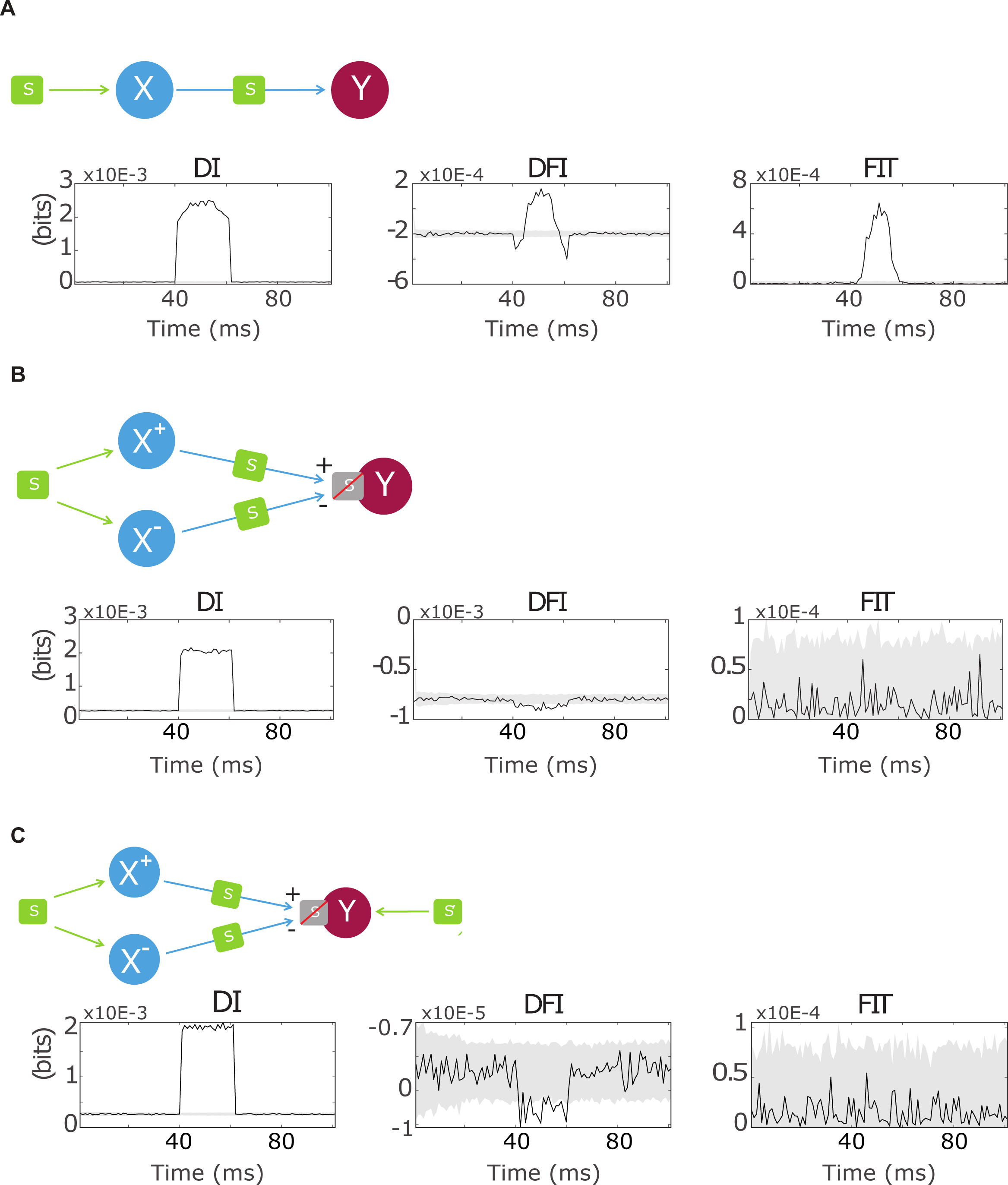
Simulations for testing Feature-specific Information Transfer. In all panels, information is sent from X to Y only in a communication window between 40 and 60 ms. Each row represents a different communication scenario. Colored upper diagrams show a schematic of the information transfer scenario considered in each row. Second, third and left columns from left display the time course of the DI, DFI and FIT, respectively. In these panel, black lines correspond to average over simulations, and grey areas correspond to confidence intervals (0.1-99.9 percentiles) of the mean computed from surrogates. **A.** Communication scenario representing feedforward transmission about the considered stimulus feature. The communication interval from 40 to 60 ms is captured by all measures as a significant increase with respect to baseline information. **B.** Communication scenario representing a transfer of information, that is not about the considered stimulus feature, from X to Y. The DI captures the presence of information, but cannot reveal its stimulus specificity. Both the DFI and FIT correctly capture the lack of stimulus-specific transfer. **C**. Communication scenario representing an information transfer from X to Y that does not convey information about S, though the information is present in X. Note that an extra source of information about a different stimulus S’ drawn randomly from the same probability distribution as S (left panel). Whereas the DI shows an increase in information transfer, both the DFI and FIT correctly detect the absence of stimulus-specific information transfer. We should note, however, that the DFI (unlike the FIT) contains significant negative values both in (B) and (C).

The first scenario (schematized in Fig. 1A, top) modeled a transfer of information from *X* to *Y* that was specifically about the feature *S* distinguishing the different possible simulated input stimuli. Since in our simulations each stimulus elicited a peak in activity in the sender X with the same temporal shape but with different stimulus-dependent peak heights, the feature *S* distinguishing the possible simulated stimuli was the height of the activity peak in sender *X*. As expected, all three measures captured the information transfer and showed a significant peak in the time period in which information was actually transferred (Fig. 1A). However, the DFI displayed negative values in the periods outside the information transfer window. These negative values are hard to interpret (Ince et al., 2015) and were not found with the FIT measure which, by definition, is non-negative (Fig. 1A).

The second scenario mimicked a transfer of information from X to Y that was feature-unspecific. This was achieved by combining the input from X to Y in such a way that the stimulus-related signal in X is canceled out when received in *Y* (Fig. 1B top, see Methods). This scenario, although admittedly contrived, is useful to test the ability of different measures to distinguish between feature-related and feature-unrelated information transfer. DI showed a significant increase in the information transfer window (Fig. 1B, left panel). This was expected, because the DI lacks the ability to discern feature-related from feature-unrelated information transfer. However, both the DFI and FIT did not show any increase during the information transfer period, correctly detecting the lack of stimulus-related information transfer (Fig. 1B, right panels).

In the third scenario (Fig. 1C), we simulated a condition in which an external confounding stimulus signal was injected to the receiving node *Y*, while the information transferred from *X* to *Y* was not about the stimulus feature (Fig. 1C, left panel). In this case, both *X* and *Y* have stimulus feature information, but the feature information in *Y* did not come from *X*. The DI indicated an information transfer between *X* and *Y*, but could not reveal if it was about the stimulus feature. However, neither the DFI nor FIT measures showed a significant peak at any time, correctly picking that the transfer from X to Y was not about the stimulus feature. We should note, however, that the DFI, as in Fig. 1B, exhibited significant negative values during the transfer period.

Overall, the simulation results indicate that the FIT measure successfully tracks genuine feature-related information transfer and, unlike the DFI, it provides non-negative estimates.

To further demonstrate that FIT reliably reflects transmission of information about the considered feature, rather than transmission of information that is not about the considered feature (for example, internal state variables), we performed (Fig. 2) more detailed numerical simulations using the first scenario (genuine transmission of feature-related information from *X* to *Y*). We varied systematically the amplitude of the noise parameter (the parameter regulating the amplitude of trial to trial, stimulus-unrelated variability) and of the stimulus signal parameter (that is, the amplitude of the stimulus-related response). The results showed that DI increases both with the level of noise and of stimulus signal (Fig. 2A,B, left and right panels). This means that DI, as predicted by the theoretical considerations, does not dissociate between noise-related and feature-related components of the information transfer. On the other hand, the FIT measure decreased with increasing noise level and increases with increasing stimulus signal amplitude (Fig. 2AB, center and right panels), thus demonstrating that the FIT measure focuses only on feature-related information transfer. The DFI (results not shown) showed a pattern similar to that of the FIT in terms of increase with signal and decrease with noise, although this pattern was less pronounced than that of FIT. In addition, the DFI could take negative values. Overall, these facts suggest that DFI focuses mainly on feature-related transfer but less precisely so than the FIT.

**Figure 2.**
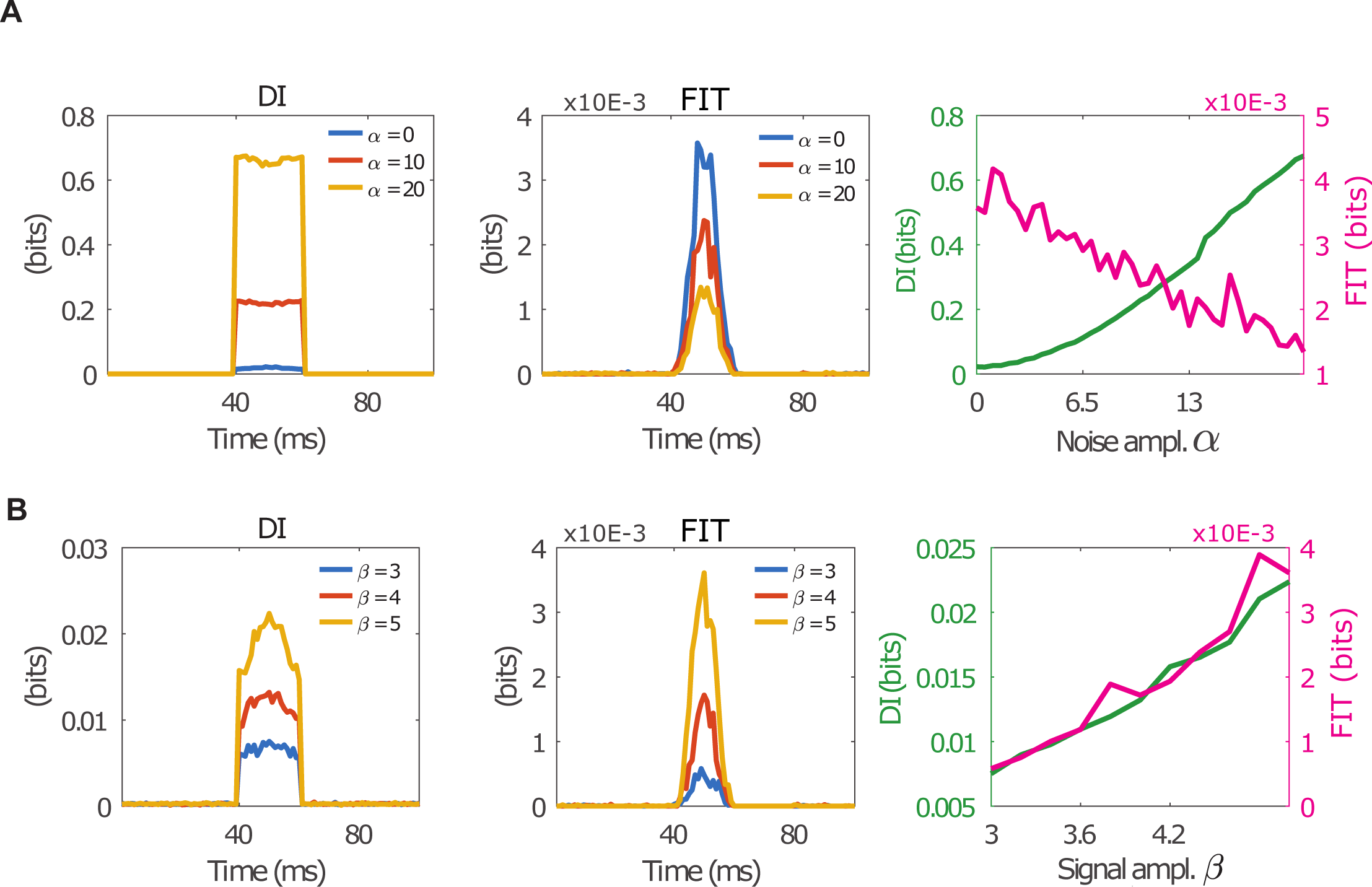
Simulation of the dependence of FIT on noise level and signal amplitude. **A.** Dependence on noise level (*α* coefficient) of the DI (left panel and right panel) and FIT (central panel and right panel). Whereas the DI increases with noise level, the FIT decreases with noise level. **B**. Dependence on stimulus amplitude (*β* coefficient) of the DI (left panel and right panel) and FIT (central panel and right panel). Both the DI and FIT increase with stimulus amplitude.

We then addressed the effectiveness of the different measures in detecting the directionality of information transfer between neural populations. We performed (Fig. 3) simulations based on the first scenario, in which by construction feature-related information was transmitted only from X to Y, but not vice versa. In all simulated cases, the information transmission period had a width of 80 ms and included a Gaussian stimulus-evoked activity bump of standard deviation 27 ms. However, we considered three different values of the delay in information transmission (5, 15 and 25 ms respectively, different rows of Fig. 3). Given that a longer delay is a clearer indication of directionality of information transfer, we expect that, for analysis algorithms, it is more difficult to determine the directionality of transfer when the delay is small. Detection of directionality was expected to be especially challenging with a delay of only 5 ms, which was much shorter than the autocorrelation in the signals (which is of the order of the width of the Gaussian stimulus-evoked activity bump shown in Fig. 3, left panels). Indeed, we found (Fig. 3) that the DI and DFI did not provide clear directional information when the delay was short with respect to signal autocorrelations (top row in Fig. 3). However, the FIT correctly detected the directionality of information transfer from *X* to *Y* in all cases (Fig. 3, right panels). Another interesting result was that the FIT was non-zero in a temporally more confined region than DI and DFI. In particular, FIT rapidly went to zero after the information about the feature carried by the sender X became smaller than the information that was already present in the past of the receiver Y. In our simulations, this happens at approximately 70 ms post-stimulus (Figure 3, right panels). This finding suggests that FIT correctly captures the fact that when this happens, information in Y cannot be imputed any more as coming from the past of X, and so communication of feature-related information from X to Y can be ruled out as unlikely. This does not happen to DI, because it does not consider the stimulus-specificity of information, and to DFI, because of its lesser sensitivity to true redundancy and uniqueness of information (Fig 3). Thus, FIT has the useful properties of being conservative when determining the temporal interval in which stimulus information is transferred, restricting the measured transfer to periods when X keeps providing new stimulus information to Y.

**Figure 3.**
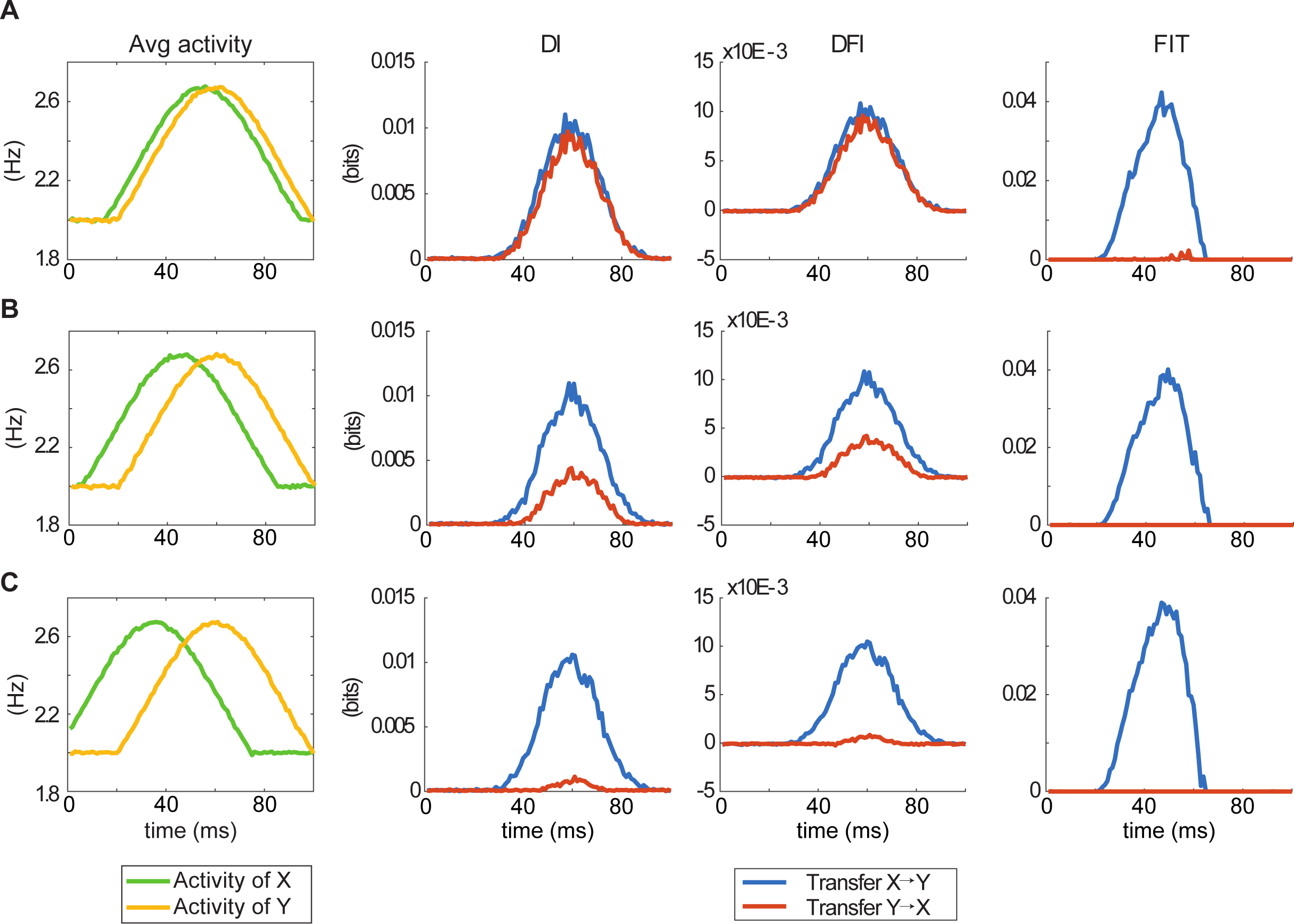
Simulation of directional-specificity and temporal precision of FIT. We simulate a scenario in which feature-specific information is transmitted only feedforward from area X to area Y. To test the directional sensitivity of the measures, we perform an information flow analysis both from X to Y (blue curves) and from Y to X (red curves). The left column shows the trial-averaged simulated activity of X (green curve) and Y (yellow curve). The DI, DFI and FTI measures are displayed in the second, third and fourth columns, respectively. **A.** Results from simulations using short distances between activity peaks (5 ms) and very large temporal overlap of neural activity in X and Y. Both the DI and DFI, but not the FIT, incorrectly detect an information flow from Y to X. **B.** Results from simulations using intermediate distances between activity peaks (15 ms) and relatively large overlap of neural activity Intermediate distance (15 ms). Similarly to the cases plotted in panel (A), both BI and DFI display a non-zero information flow from Y to X, whereas the FIT correctly detects information only from X to Y. **C.** Results from simulations using a longer delay (25 ms) of information transmission from X to Y, and thus a only moderate temporal overlap bet. In this case all three measures detect correctly directionality of information flow.

To further investigate the robustness of FIT to noise, we performed additional simulations using the first scenario and tested noise levels at the limit of detectability for the DI and DFI measures (Suppl. Fig. 1A). In the high-noise condition, we added a sine wave with a period of 200 ms and random phase at each trial to both signals *X* and *Y*. The low-noise condition had a constant value added to both signals, thus decreasing the relative difference between the baseline and the peak. Indeed, the results showed that, whereas the DI and DFI could not capture the information transfer significantly from the data (Suppl. Fig. 1B, second and third columns), the FIT was more robust to noise (Suppl. Fig. 1B, fourth column).

Then, we investigated how the different measures behave when X and Y display synergistic feature information coding. Indeed, the mathematical decomposition of the DFI in different terms of the PID framework (Williams and Beer, 2010) shows that the information redundancy measure shown by the DFI contains synergistic terms, which may lead to negative values of the measure (see Supplementary Methods, as well as (Ince et al., 2015), for more details about this problem). We thus designed a scenario (Suppl. Fig. 1C) in which the correlations between X and Y were modulated by the stimulus feature. In such case, use of the information breakdown (Pola et al., 2003) to quantify the impact of stimulus-independent and stimulus dependent correlations, shows that the stimulus-dependent correlations lead to synergistic coding of information (quantified in the *Icor-dep* term of Supplemental Figure 1C), because the feature can be decoded from the level of correlation between X and Y (Nigam et al., 2019; Pola et al., 2003). As expected, in this case the DFI yielded significant negative values, which are difficult to interpret in terms of stimulus information transfer (Ince et al., 2015), while both DI and FIT displayed a significant positive value during the transfer period (Suppl. Fig. 1C). Therefore, we conclude that the FIT successfully identified stimulus-specific information transfer even in the presence of strong synergistic effects. As a final control analysis, we investigated the susceptibility of the different measures to the limited sampling bias, which is known to affect the estimation of information theoretical measures (Panzeri et al., 2007; Panzeri and Treves, 1996). We tested the behavior of the DI, DFI and FIT on all three scenarios with varying number of data samples (Suppl. Fig. 2). The results showed that FIT, like all other information theoretic measures, suffers from an estimation bias that decreases with the number of samples (Suppl. Fig. 2). This bias can, however, be corrected using the quadratic extrapolation procedure introduced in (Panzeri et al., 2007; Strong et al., 1998). Using this bias correction, we showed that it is possible to obtain reliable values of FIT with approximately a tenth of the number of samples with respect to when not using any bias correction. We, however, did not correct for limited sampling in the main figures of this paper, because, under the conditions considered here, we found that the bias correction did not change any of the presented results. This correction, however, could be of practical benefit in conditions of under-sampling more severe than those of the datasets analyzed here, and thus its use may be of use for some data analyses.

### Tracking feature-related information flow from neurophysiological data

To investigate the ability of the FIT to track feature-related information flow from brain signals, we analyzed three sets of neurophysiological data exemplifying experimental setups encountered in cognitive and system neuroscience: non-invasive scalp-level electroencephalographic (EEG) signals and cortical-level high-gamma activity (HGA) estimates from magnetoencephalographic (MEG) data from human participants, and intracranial multi-unit activity (MUA) signals recorded in rodents.

#### Information transfer during face detection (human EEG)

Here, we tested whether the FIT could detect feature-related information flow between brain hemispheres using an EEG dataset collected from human participants performing a face detection task (Ince et al., 2016; Rousselet et al., 2014). In this task, subjects were asked to detect the presence of either a face or a random texture while the image was covered by a bubble mask randomly generated in each trial. We used FIT and other measures to investigate both the feature selectivity and the direction-specificity of the information flow across hemispheres.

Previous work on face detection in humans has shown that the first EEG component that preferentially responds to faces is the N170 (Bentin et al., 1996). Encoding of face information in the EEG starts with the encoding of the visibility in the masked image of the eye contralateral to the considered hemisphere, on the downward slope of the N170 (~140 ms post-stimulus, as shown in Fig 4A, which displays the mutual information about the left eye visibility carried by the right hemisphere EEG). Left eye visibility is an image feature highly relevant for face discrimination at the behavioral level (Ince et al., 2016). Encoding of information about the ipsilateral eye, another behaviorally relevant face feature, appears later in the EEG, at approximately 180 ms (see (Ince et al., 2016), and Fig 4A which shows the mutual information about the left eye visibility carried by the left hemisphere EEG).

**Figure 4.**
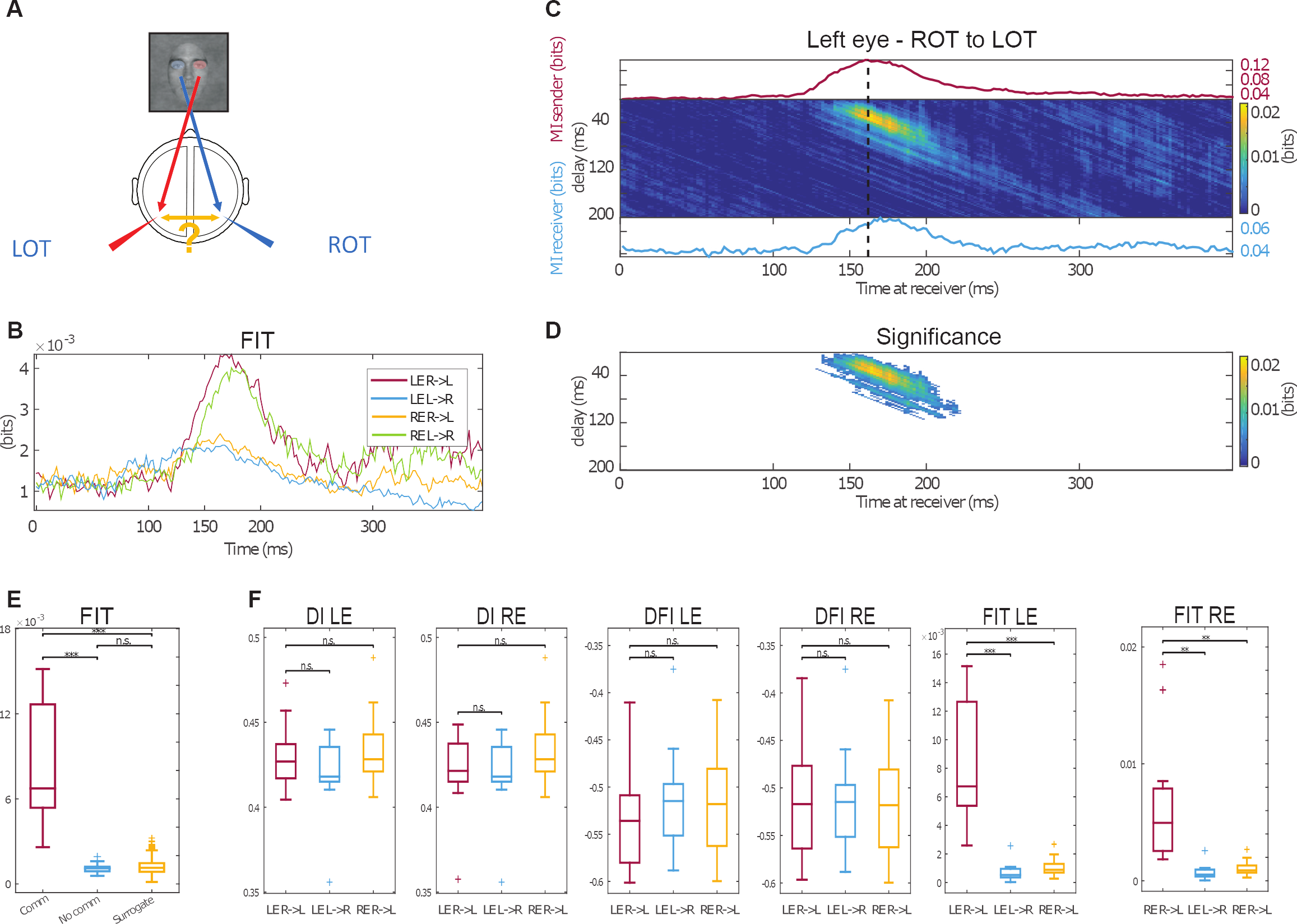
Inter-hemispheric eye-specific information transfer in face detection using human EEG. **A.** Schematic of the investigated information flow patterns. We expect that information about the presence of an eye in the image is initially encoded in the hemisphere contralateral to the eye, and we investigate whether this information is then transferred to the other hemisphere. LOT and ROT refer to the Left and Right Occipito-Temporal EEG sensors, respectively. **B.** Image plot the values of the FIT, as function of the two parameters communication delay and post-stimulus time, capturing the transfer of information about visibility of left eye from the right to the left hemisphere. Line plots above and below the image report the value of the mutual information between the given occipito-temporal sensor (either sending or receiving one) and visibility of the left eye. **C.** FIT values averaged across delays for all combinations of transfer direction and eye. **D** Image plot (as function of the parameters communication delay and post-stimulus time) of the values of the FIT (about the visibility of left eye, from the right to the left hemisphere) that are significant after a cluster-based nonparametric permutation test (see Methods). The (time, delay) parameter region with significant values of FIT was defined to be the communication window. **E.** FIT values averaged within the communication window (indicated as comm) compared to the values averaged over a (time, delay) cluster of the same shape and size, but positioned in a pre-stimulus with a long delay and no overlap in which communication was not significant and not possible (indicated as “no comm”), and surrogate values obtained by random shuffling of the stimulus values (left eye visibility). **F.** Comparison between a given contra to ipsilateral transfer (L – left, R - right) and the two ipsi-to contra-lateral transfers for DI, DFI and FIT and both eyes (LE and RE). In Panels B-D, we plot and compute the statistical significance of the average across subjects of the information values. In panels E,F boxplots represent distributions of information values over subjects.

A question highly relevant to understand the brain mechanisms of face recognition is whether the later information about the ipsilateral eye is computed locally or not. A recent study (Ince et al., 2016) hypothesized that eye-related information first appears in the hemisphere contralateral with respect to the position of the eye, and then flows from the contralateral to the ipsilateral hemisphere (Fig. 4A). Ince and colleagues (Ince et al., 2016) corroborated this hypothesis by showing that information about a given eye encoded ipsilaterally later on, was partly redundant with the information about the same eye encoded contralaterally at earlier times. Despite this evidence in support of a direction-specific cross-hemisphere transfer of contralateral eye information, this previous study did not demonstrate that information about a given eye that flowed from the contralateral to the ipsilateral hemisphere was indeed absent ipsilaterally at earlier times.

To investigate this hypothesis and demonstrate the presence of feature information transfer, we computed transmission of eye-related information across hemispheres using FIT. We first considered transmission of information about the presence of the left eye in the face image. The FIT values clearly showed a transfer of left-eye-related information from the right to the left hemisphere (Fig. 4B). Cluster analysis of the FIT values as a function of post-stimulus time and delay of transmission identified a “communication window” of significant increase of left-eye FIT in a time interval between approximately 150 to 190 ms after stimulus-onset and transfer delay of approximately 10 to 50 ms (Fig. 4C, D). Left-eye related FIT information from right to left hemisphere averaged in the significant time-delay region (referred to as “comm” in Fig. 4E) was significantly higher than the FIT values averaged over a cluster of the same shape and size, but positioned elsewhere on the timeline with no overlap (“no comm” in Fig. 4E) and to the null-hypothesis FIT values obtained by randomly shuffling left-eye visibility values. Thus, the FIT robustly determined the communication window for right-to-left transfer of left-eye information with high temporal precision.

To gain further insight about the directionality and feature-specificity of the information transfer across hemispheres, we computed FIT for all four possible transfer directions and positions of the eye on the image (i.e., from ipsi to contralateral and vice versa, and left and right eye, respectively). For these analyses, all presented in Fig. 4F, we averaged information values in the time-delay region displaying significant FIT (this region, plotted for left eye information in Fig. 4B, included those in which DFI and DI were significant, as FIT is more temporally selective). We found that left-eye related FIT was present in the right-to-left hemisphere direction, but not in the opposite direction. The FIT in the right-to-left hemisphere direction was present only about left-eye information, but not about right eye information (Fig. 4F). Similarly, right-eye related FIT was present only in the left-to-right direction (not in the opposite one), and that left-to-right hemisphere FIT was present only about right-eye (but not left-eye) information (Fig. 4F). Together, the use of FIT allowed us to reveal temporally localized transfer of visual information that was feature-selective (about only the contralateral but not the ipsilateral eye) and direction-specific.

When using either DI and DFI, no difference was found between directions of transfer (left-to-right vs right-to-left hemisphere, see Fig. 4F) both for right and left eye. For DFI, when considering a given direction of transfer (left-to-right or right-to-left hemisphere, see Fig. 4F), no difference was found between left-eye-related and right-eye-related transmission. Since DI is not feature-dependent, of course it could not detect difference between left-eye-related and right-eye-related transmission. Thus, while the feature selectivity and direction-specificity of inter-hemispheric information transfer was expected from empirical considerations, neither the DI or DFI could show this effect, whereas FIT could.

#### Thalamocortical feature-selective information transfer during multisensory stimulation (rodent MUA)

To further test the FIT on real neurophysiological data, we analyzed the thalamocortical information flow along the somatosensory and visual pathways. As above, we aim at testing the feature selectivity and direction-specificity of the FIT. In the neural systems under consideration, there is a strong expectation that (i) information about basic sensory features in the receptive field flows primarily feedforward (from thalamus to cortex) rather than in the feedback direction, (ii) that the feedforward somatosensory pathway transmits more information about somatosensory features than visual features, and that (iii) the feedforward visual pathway transmits more information about visual features than somatosensory features.

In this dataset, spiking multi-unit activity (MUA) was simultaneously recorded in the primary visual cortex (V1), the first-order visual thalamus (the lateral geniculate nucleus, LGN), the primary somatosensory cortex (S1) and the first-order somatosensory thalamus (the ventral posteromedial nucleus, VPM) of anaesthetized rats (Fig. 5A). Neural activity in these regions was recorded in response to receptive field stimulation that was either unimodal visual, unimodal somatosensory or bimodal (simultaneous visual and tactile). From these three stimuli, we constructed two stimulus feature sets. The first stimulus set consisted of the unimodal visual and the bimodal visual-tactile stimulus. We call it the somatosensory-discriminative set, because their two stimuli are discriminated by the presence or absence of tactile stimulation. The second stimulus set, that we call the visual-discriminative set, was made of the unimodal tactile and the bimodal visual-tactile stimulus. By computing FIT about either the somatosensory-discriminative or the visual-discriminative stimulus set, we could measure the transfer of information about either somatosensory or visual features.

**Figure 5.**
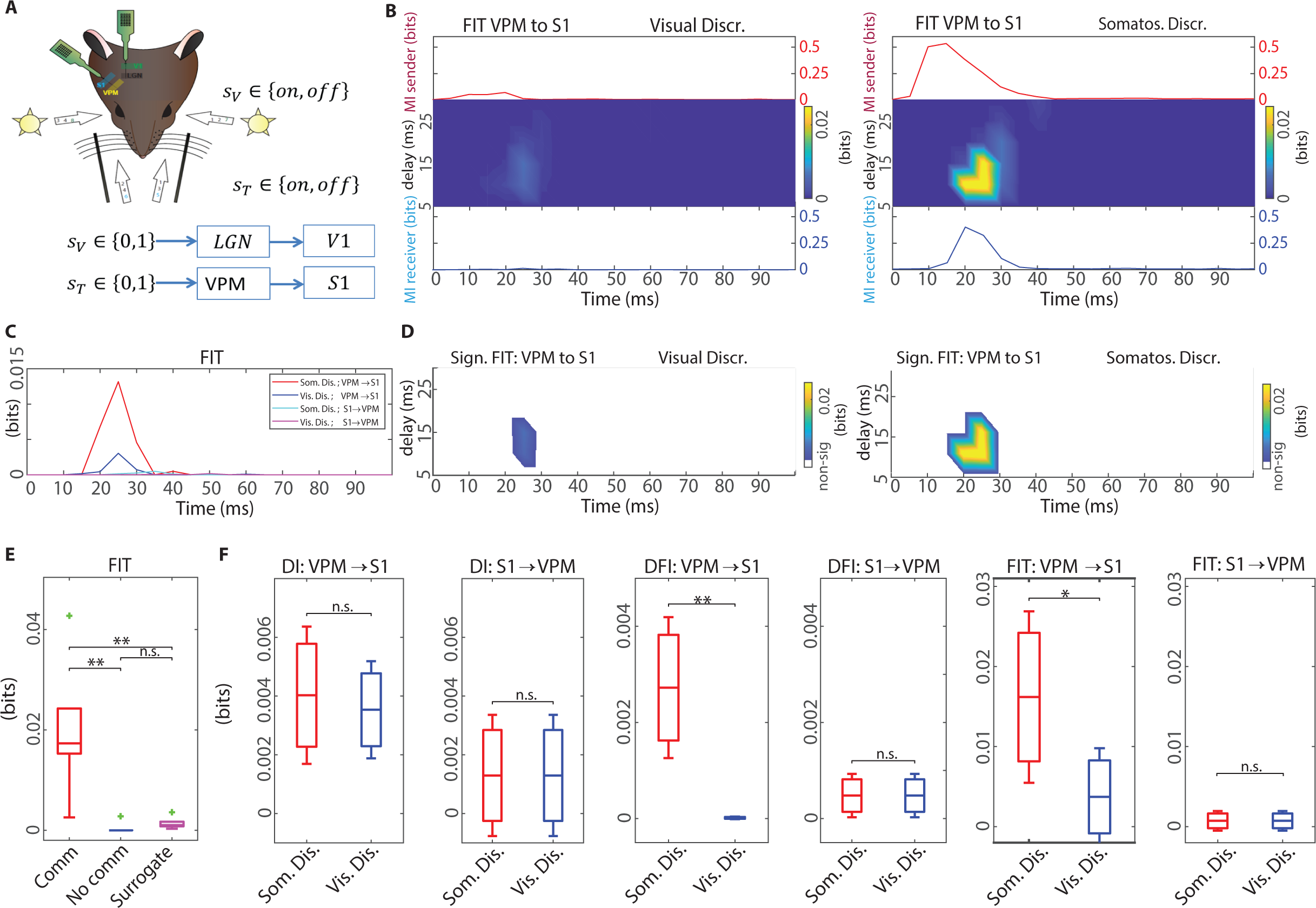
Sensory-related information transfer in the somatosensory system (rat MUA). **A.** Schematic of the experiment. MUA was recorded from the visual and somatosensory territories of the thalamus and associated primary cortical regions in rats. We considered only sensory stimulation contralateral to the recorded regions. Visual information flows from the eye to the primary visual cortex (V1) through the Lateral Geniculate Nucleus (LGN) of the thalamus and somatosensory information flows from the whiskers to the primary somatosensory cortex (S1) through the Ventral Posteromedial nucleus (VPM) of the thalamus. **B.** FIT from VPM to S1. The image plots shows the values of FIT (mean across subjects) for each value of delay and post-stimulus time. Line plots above and below show Mutual Information (MI) between the presented stimulus and the recorded MUA in the VPM and SI, respectively. The left panel reports values of information and FIT about the visual discriminative stimulus set, whereas the right panels report values of information and FIT about the somatosensory-discriminative stimulus set. Information flow from somatosensory thalamus to somatosensory cortex is only about the somatosensory discriminative feature. **C.** FIT values averaged across delays for all combinations of transfer direction and stimulation type. **D.** Image plot (as function of the parameters communication delay and post-stimulus time) of the values of the FIT that are significant (p< 0.05) after a cluster-based nonparametric permutation test (see Methods). The (time, delay) region with significant values of FIT was defined to be the communication window. The left panel reports FIT about the visual discriminative stimulus set, whereas the right panel reports FIT about the somatosensory-discriminative stimulus set. **E.** Comparison of FIT values averaged within the (time, delay) communication cluster (indicated as “comm”) with the values obtained averaging FIT within a cluster of the same shape and size, but positioned elsewhere in a prestimulus time window and long delay region in which communication was neither significant nor possible (“no comm”), and surrogate FIT values obtained by random shuffling of the stimulus values. **F.** Comparisons between all 4 combinations of transfer direction (from thalamus to cortex and vice-versa) and conditions (Somatosensory and visually discriminative), for all the three measures. In Panels B-D, we plot and compute the statistical significance of the average across subjects of the information values. In panels E,F boxplots represent distributions of information values over subjects.

We first considered the somatosensory pathway (VPM and S1) and the somatosensory-discriminative stimulus set. Somatosensory-discriminative information in neural activity was present in the 5-30 ms and 15-30 ms post-stimulus intervals in VPM and S1, respectively (Fig. 5B).

Cluster analysis (Maris and Oostenveld, 2007) of the somatosensory-discriminative FIT values from VPM to S1, computed as function of post-stimulus time and delay of transmission, identified a communication window of somatosensory information transmission from VPM to S1 in a time interval between approximately 15 to 30 ms after stimulus-onset and transfer delay of approximately 10-15 ms (Fig. 5B, D). FIT values were significantly higher in this communication window than those obtained at different time points and with respect to surrogates (Fig. 5E).

Similar results were found, using FIT, in the visual system for transmission of visually-discriminative information (Suppl. Fig. 3). In this case, the cluster analysis (Maris and Oostenveld, 2007) of FIT values identified a communication window of significant visually-discriminative information transmission from LGN to V1 in a time interval between approximately 50 to 70 ms after stimulus-onset and transfer delay of approximately 10-20 ms (Suppl. Fig. 3B, D).

Thus, the FIT measure determined the window for feedforward communication of sensory information from thalamus to cortex with high temporal precision and robustness.

To test the directional specificity and feature selectivity of the information transfer within the thalamo-cortical pathway, we computed FIT for both feedforward (thalamus to cortex) and feedback (cortex to thalamus) direction, for both the somatosensory- and the visually-discriminative stimulus set. Results for the somatosensory system are reported in Fig. 5B, F. We found that, for VPM to S1 communication, somatosensory-discriminative FIT values were significantly larger than the visually-discriminative ones (Fig. 5F; p = 0.032). Moreover, somatosensory-discriminative FIT values were larger in the feedforward than in the feedback direction (Fig. 5F; p = 0.0056). Results for the visual system are reported in Suppl. Fig S3. We found that, for LGN to V1 communication, visually-discriminative FIT values were significantly larger than the somatosensory-discriminative ones (Suppl. Fig. 3F; p = 0.019). Moreover, visually-discriminative FIT values were larger in the feedforward than in the feedback direction (Suppl. Fig. 3F; p = 0.0087).

The FIT results imply that the sensory information in the thalamocortical system is primarily feedforward and that predominantly visual information flows through the visual system and predominantly somatosensory information flows through the somatosensory system. While these results may sound trivial, they are difficult to obtain, even from the same dataset with established or state-of-the-art methods used to estimate directed information flow. This is apparent when considering the results obtained with DI or DFI. In the somatosensory dataset, modulations in feedforward and feedback DI were similar (p=0.68 and p = 0.052, respectively) between the visually-discriminative and somatosensory discriminative datasets (Fig. 5F, top-left panel). Furthermore, the difference between feed-forward and feedback somatosensory-discriminative DI was barely significant (p=0.058). Comparable results were found when examining the values of DI in the visual system (Suppl. Fig. 3). The DFI showed more direction-specificity and feature-selectivity than the DI. However, unlike the FIT, the DFI was able to capture the feedforward feature-specificity only for the somatosensory pathway (p=0.0011 for VPM->S1, Fig. 6F, top-right panels), but not for the visual pathway (p=0.125 for LGN->V1, p=0.321 for V1->LGN) (Suppl. Fig. 3F, top-right panels).

**Figure 6.**
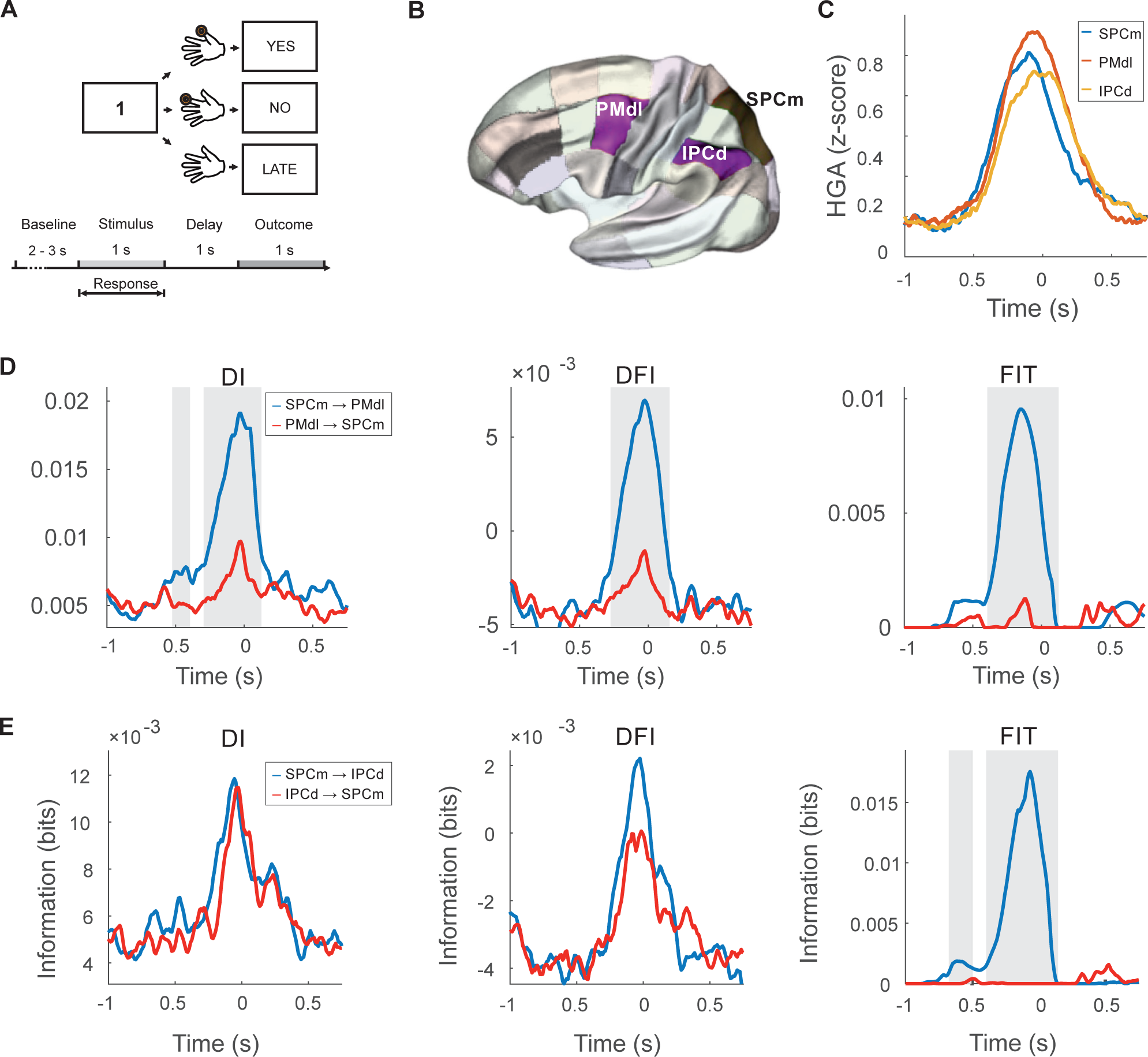
Information transfer in high-gamma activity (HGA) during a visuomotor mapping (human MEG). **A.** Schematic of the arbitrary visuomotor mapping task. **B.** The anatomical regions-of-interest used in the current study (the dorsal Inferior Parietal Cortex, IPCd; the medial superior parietal cortex, SPCm; and the dorsolateral premotor cortex, PMdl) are shown in color superimposed to a gray-shaded *MarsAtlas* cortical parcellation scheme. **C.** Mean HGA averaged across trials, sessions and participants for the three regions-of-interest. HGA is expressed as z-score with respect to baseline activity (from −0.5 to −0.1s) prior to stimulus onset. **D.** Information flow between the SPCm and the PMdl estimated using the DI (left panel), DFI (central panel) and FIT (right panel). The blue and read lines shows flow measured from SPCm and PMdl, and from PMdl to SPCm, respectively **E.** Information flow between the SPCm and the IPCd estimated using the DI (left panel), DFI (central panel) and FIT (right panel). The blue and read lines shows flow measured from SPCm and IPCd, and from IPCd to SPCm, respectively.

Taken together, these results highlight the power of the FIT in revealing credible temporally localized transfers of information that are feature-selective and direction-specific.

#### Information transfer in human visuomotor network (MEG)

In the third set of analyses, we used the FIT to measure transfer of information signaling engagement in visuomotor transformation within the human visuomotor network. We analyzed magnetoencephalographic (MEG) data recorded while participants were asked to perform finger movements associated to digit numbers presented on a screen. The appearance of digit “1” instructed the execution of the thumb, “2” for the index finger, “3” for the middle finger, and so on (Fig. 6A, see (Brovelli et al., 2015)). We analyzed single-trial source-level high-gamma activity (HGA; defined as the instantaneous power of the 60-120Hz frequency band). HGA has been shown to capture local spiking activity (Ray and Maunsell, 2011), and it can be used for mapping cognitive-related areas (Cheyne and Ferrari, 2013; Crone et al., 2006; Darvas et al., 2010; Jerbi et al., 2009; Lachaux et al., 2012) and functional connectivity networks in humans (Brovelli et al., 2017, 2015).

Previous work has shown that the medial superior parietal cortex (SPCm) plays a key role in the visuomotor-related functional network (Brovelli et al., 2017). The SPCm is identified to be at the top of the cortical hierarchy by two measures, eigenvector centrality and betweenness centrality, which quantify the propensity of a brain area to participate in multiple sub-networks. In particular, the SPCm was found to participate in both a visuo-parietal network early during the processing of visual information and during the planning of visuomotor associations with the dorsolateral premotor cortex (PMdl). Granger causality analyses revealed a directional influence from the superior parietal lobe to the dorsal premotor areas (Brovelli et al., 2015). However, other areas, such as the dorsal Inferior Parietal Cortex (IPCd), responded almost simultaneously with respect to SPCm (Fig. 6C), and it was difficult to identify the directionality of the information flow between SPCm and IPCd. In Fig. 6B we plot the regions-of-interest considered in our analyses using the MarsAtlas parcellation scheme (see Methods and (Auzias et al., 2016) for more details).

We subjected single-trial HGA to FIT, DI, and DFI analyses to assess the dynamics of transfer of information within the visuomotor network. We constructed a stimulus feature set that was made of the baseline condition (no task) vs the visuomotor execution. By computing FIT or DFI about this feature stimulus set, we could measure transfer of information that signals engagement in visuomotor mapping. For comparison, DI was computed using data taken both during baseline and task condition. When considering the information transfer between SPCm and PMdl (Fig. 6D), we found that all measures showed a time-resolved directional pattern (higher directionality from the SPCm towards the PMdl), supporting previous static (i.e., non-time-resolved) Granger Causality results (Brovelli et al., 2015). This result confirms, with more general information theoretic methods and temporal dynamics information, that the interaction between the superior parietal lobe and the dorsolateral premotor cortex dynamically occurs over a timescale of tens to hundreds on milliseconds during a time interval prior to motor output. Moreover, the finding that both FIT and DFI are directional demonstrates that the transfer of information from SPCm towards the PMdl specifically carries information signaling the execution of a visuomotor transformation, something that could not be established by Granger Causality or DI alone.

We then analyzed the information flow between the SPCm and IPCd (Fig. 6E). We found that the FIT displayed a clear and significant selectivity of the directionality of information flow. The flow was absent in the direction from IPCd to SPCm, but was strong, and well time-localized prior to finger movement, in the direction from SPCm to IPCd (Fig. 6E). DFI and DI instead did not reveal a significant direction selectivity of information flow between these two areas.

Overall, these results suggest that the FIT may be a powerful tool for mapping, in the human brain using MEG data, feature-selective and direction-specific information transfer at a high temporal resolution, essential requisites for attempting to track cortico-cortical directional interactions.

## Discussion

Here, we developed and validated a novel information theoretic measure that quantifies feature-specific information transfer (FIT) across two neural signals, *X* (the sender) and *Y* (the receiver). This measure is designed to capture the transfer of information about the considered stimulus feature *S*. It is based on the Partial Information Decomposition formalism (Williams and Beer, 2010) and on the notions of redundancy and uniqueness of information. The rationale is that, if there is transfer of information about a feature *S* from *X* to *Y*, then information about *S* is shared (redundant) between the past of *X* and present of *Y*, while being absent in the past of *Y*. In intuitive terms, our measure considers the information about *S* encoded by the neural signal *Y* at the present time, and takes from it only that part of information that is shared with information about *S* previously encoded in *X* and removes from it the information about s that was present in the past activity of *Y*.

We performed numerical simulations to test the reliability of the FIT measure to capture feature-specific information transfer in different communication scenarios, and we compared its effectiveness against two already available measures. The first is DI, the most established information theoretic version of Granger Causality, designed to capture the transfer of information without regard to the specific features about which the transmitted information is about. The second is DFI, a recent attempt to capture feature-related transfer that reasoned along similar intuitive terms, but did not use the PIDs to identify the redundancies and uniqueness of information representations. In short, the simulations confirmed the properties of each measure that was expected from the above theoretical reasoning. DI was very effective at detecting transfer, but it did not identify the feature specificity of this transfer. FIT was very effective at detecting feature-relatedness and feature-specificity of the information transfer. DFI had some effectiveness for detecting feature-related transfer, but was much less effective and harder to interpret than FIT. Simulations also revealed an excellent temporal sensitivity of FIT in detecting genuine transfer in the correct post-stimulus time ranges and communication delays. The ability to detect transfer at short delays (shorter than the autocorrelations of the signals) was instrumental in ultimately leading FIT to have an excellent direction-sensitivity.

The excellent properties of FIT in terms of feature-selectivity, direction sensitivity and temporal resolution found in simulated data were all strongly confirmed in the analysis of real data. In human EEG data recorded during a face detection task, FIT was able to reveal the feature specificity and directionality of interhemispheric information flow, determining that in this task interhemispheric communication transmits information primarily about the visibility of the contralateral eye in the direction from the contralateral to the ipsilateral hemisphere. In MUA data recorded from the thalamocortical sensory pathway, FIT revealed that modality-specific information (somatosensory information for the somatosensory pathway, visual information for the visual thalamocortical pathway) predominantly flows in a feedforward manner (from thalamus to cortex). In MEG data, FIT could reveal a clear directionality in the information flow in visuomotor association tasks. In all cases, FIT showed a sharp temporal sensitivity in individuating the window in which communication takes places. We believe that this fine temporal and directional sensitivity arises from the power of the PID approach to consider only unique information that did not appear in the past.

The DFI could detect expected feature specificity from neurophysiological data only in a subset of cases (a somatosensory information flow in the somatosensory thalamocortical cortical pathway, and a fronto-parietal information flow from superior parietal to premotor area in MEG data), but it could not detect feature specificity in other cases (feature specificity of the interhemispheric information flow in EEG face detection data, flow of visual information through the thalamocortical pathway in MUA data). DFI failed to detect direction-specificity in all real data applications apart from one single case (transfer of information from superior parietal to premotor area in visuomotor associations with MEG data). Overall, our data support that FIT should be used as a measure of feature-specific transfer, as FIT showed a much higher reliability, interpretability and sensitivity with respect to other existing measures.

Neurophysiological data analyses using the DI measure confirmed its very good performance in terms of detecting feature-unrelated information transfer. On the other hand, the DI, by construction, was unable to determine whether or not the transfer related to specific features. In general, DI applied to real data confirmed the simulation results of a poorer temporal sensitivity and a lesser direction-sensitivity with respect to FIT. We believe that such lower temporal and directional sensitivity arises from the fact that DI discounts past information simply by conditioning in past activity, rather than identifying and removing information that was common between the present and the past, as done with the PID in the FIT definition. This suggests that also feature-unrelated measure of information, such as the DI, could be improved in future work by using PID, rather than conditioning, to identify only information in present activity that was unique and novel with respect to information that had been present in past activity, a key requirement to pinpoint the receipt of new information.

The success of the FIT in detecting feature-specific direction-selective transfer of information at high resolution underlies the potential power (for mapping information flow in neural circuits) of the idea of mapping transfer of information about a feature *S* from *X* to *Y* by identifying information about *S* is shared (redundant) between the past of *X* and present of *Y*, while being absent in the past of *Y*. Our new definition reached this success by extending in crucial ways previous attempts to map feature-specific transfer. In previous work that developed the DFI (Ince et al., 2015), the idea of identifying feature information in Y that was at the same time redundant with that in the past of X and unique with respect to that in the past of Y was implemented. However, the DFI’s use of an information redundancy measure that included also synergistic information, and the use of conditioning to identify information that was novel with respect to test past led to major problems in making the measure work, as demonstrated by our simulations. Other work proposed to map feature-selective transfer from X to Y using the PID concept of intersection information (Panzeri et al., 2017; Pica et al., 2019, 2017), mapping feature-related information that is shared between the past of *X* and the present of *Y*. This earlier preliminary work, however, did not derive how to discount feature information present in the past of *Y*. This ultimately makes this earlier method unable to fulfill a key notion to map genuine information transfer, i.e., the Wiener-Granger principle (Granger, 1969; Wiener, 1956). This earlier method would fail in cases in which the same feature-related information was already present in the past of Y well before it was present in the past of *X*, and when thus genuine feature-specific information transfer from X to Y would not have been credible. The advances we made in the present paper were thus not incremental, but fundamental for the FIT to reach strong performance and credibility in mapping feature-related transfer.

Recent technological progress is beginning to allow neuroscientists to both image and perturb with single-cell resolution and high temporal precision potentially groups of individual neurons (Emiliani et al., 2015). The feature selectivity of FIT and its ability to pinpoint the time and delay at which information transfer happens seems ideally suited to accompany the imaging-and-perturbation experiments that test causally information flow during cognitive tasks involving perceptual discrimination of a certain stimulus feature. For example, knowledge, obtained by FIT analysis of imaging data, could help experimenters formulate quantitative hypotheses about which populations need to be perturbed at each instant of time to perturb information flow within the brain to maximally alter behavioral task performance. Thus, an important line of future research is to understand how best to combine statistical methods such as FIT and perturbation methods to map causal feature-specific information transfer during cognitive tasks (Panzeri et al., 2017).

Despite its empirical success, future work may increase the range of applicability and rigor of FIT. First, a potential limitation of the FIT measure resides in the appropriate quantification of unique, redundant and synergistic information. In the PID, this identification of these three types of information can be done when providing a definition of redundant information. Here (see Materials and Methods), we used a particular definition of redundant information that has been presented in the literature (Williams and Beer, 2010). However, there is currently not a full agreement on what the ideal definition of redundant information could be, and the determination of the best possible measures of shared and unique information is a current hot topic of mathematical research (Barrett, 2015; Bertschinger et al., 2014; Griffith and Koch, 2012; Harder et al., 2013; Ince, 2017; Wibral et al., 2017). Performance of our FIT measures could be potentially further improved by including further advances in PID theory. Second, our current formulation of FIT is based on direct sampling of the time dependent probability of activation of neural signals *X* and *Y* as function of time and feature values. Sampling these probabilities directly is data intensive and, in practice, limits the application of FIT to univariate definitions of activity of *X* and *Y*. To apply FIT to other interesting cases that require multivariate quantification of *X* and *Y*, such as considering the transfer of information between two population codes made by many neurons in different areas, it would be important to understand how best to couple the FIT to either parametric (Sheikhattar et al., 2018) or non-parametric (Safaai et al., 2018) methods to quantify robustly neural population probabilities.

To conclude, the results from numerical simulations and the analysis of human and rodent neurophysiological data showed that the FIT provides high discriminability of information transfer both in time and across directions of information flow. Altogether our work suggests that the FIT measure has potential to uncover previously hidden feature-specific information transfer from neural data and provide a better understanding of brain communication.

## Materials and Methods

### Definition and calculation of Directed Information (DI)

The amount of transfer of information between two random variables *X* and *Y*, is usually quantified according to the Wiener-Granger principle using measures such as Transfer Entropy (Schreiber, 2000) and Directed Information (Massey, 1990), which are intimately related (Amblard and Michel, 2011). In our work, we will use the term Directed Information (DI), for the estimate

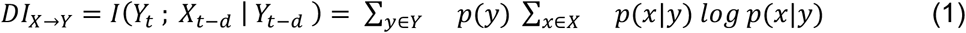

where *Y*_*t*_ is the value of *Y* at time *t*, whereas *X*_*d*−*d*_ and *Y*_*d*−*d*_ are the values of *X* and *Y* at some preceding point in time *t* − *d* (where *d* stands for delay), respectively. In Eq. (1), *I* denotes Shannon’s mutual information. For completeness, we remind that the mutual information between two variables X and Y is defined as

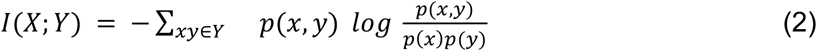

where p(x,y) is the joint probability of observing x and y and p(x), p(y) are the marginal probabilities.

DI quantifies the amount of information transferred from the past of *X* to the present of *Y*, which is not contained in the past of *Y*. DI is a model-free measure of causal information transfer in the Wiener-Granger sense, and it quantifies the amount of information between two random variables.

### Derivation of Feature-specific Information Transfer (FIT)

Here we derive mathematically the FIT measure. The aim of our work was to develop a novel information theoretical metric that quantifies the amount of information transferred from *X* to *Y* about an experimental variable *S*. Such experimental feature may represent any experimental (dependent or independent) variable under investigation (e.g., stimulus type, experimental condition, etc.). We reasoned that three conditions must be met in order for feature-specific information transfer to occur between two random processes. First, the sender *X* should encode some information about *S* at some time point in the past. Second, the receiver *Y* should encode at least part of that very same information later in time. This means that the information about S available in the present of Y should be redundant with that present in the past of X. Third, the information about S carried by the present of *Y* should be absent in the past of *Y*(otherwise the transfer would not be the origin of the information). To summarize, in order for transfer of information about *S* to occur from *X* to *Y*, the following statements should be satisfied:

1. The past of *X* at time *t* − *d* should carry information about *S*
2. The present of *Y* at time *t* should carry information about *S*
3. The past of *X* and present of *Y* should share information about *S* that is not already present in the past of *Y*

Quantifying whether or not predicate 1 and 2 are satisfied can be easily done with Shannon Information theory. The first predicate is satisfied if the mutual information between *S* and the past of *X*, namely *I*(*S*; *X*_*t*−*d*_), is positive, and the second is verified if the mutual information between *S* and the present of *Y* is positive. The third predicate needs a new measure that goes beyond Shannon information theory to be addressed. This measure is what we call FIT, and is derived as follows.

**Figure 7.**
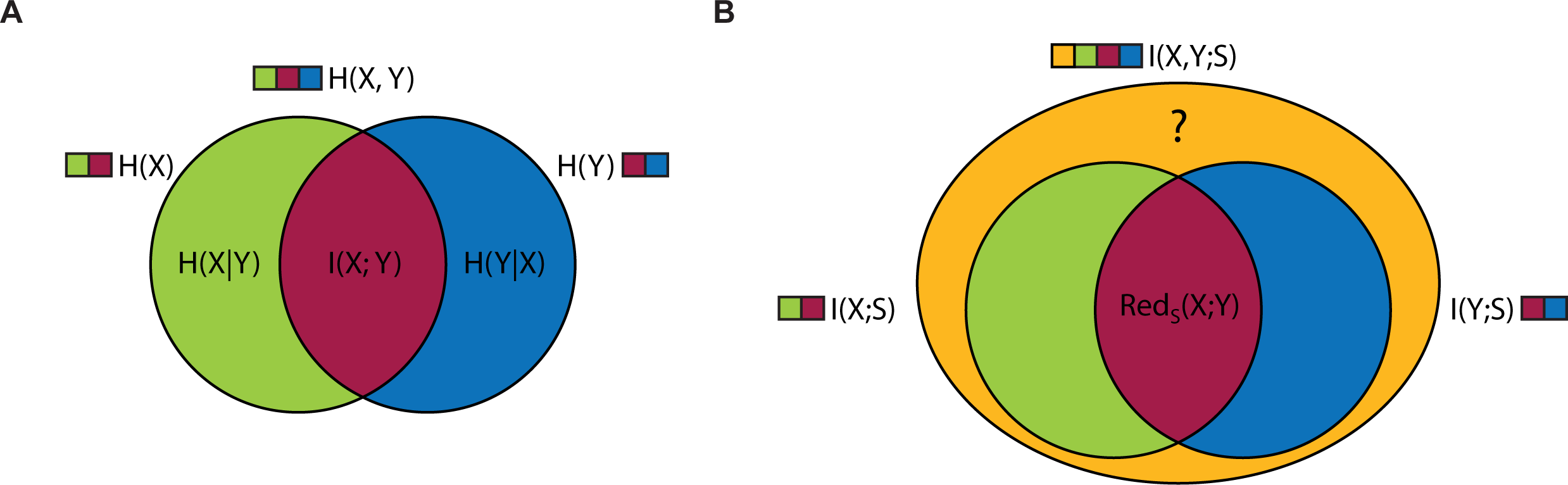
Scheme of the partial information decomposition used for FIT. **A.** Sketch of an information decomposition of interaction of two variables. *H* indicates the entropy and *I* the mutual information. The decomposition consists of three distinct parts, two representing information contributed by the variables independently (conditional entropies) (green and blue) and one that represent information that is provided by both of them (mutual - also called redundant or shared - information) (red). Based on the diagram it is possible to deduce that, to compute the mutual information, it is possible to add information contributed by both variables *H(X)* and *H(Y)* (1 green, 1 blue, 2 red parts) and subtract the total information *H(X, Y)* (1 green, 1 blue and 1 red part). **B.** Information decomposition of information interaction of two variables (*X* and *Y*) with a third one (*S*). The decomposition follows similar logic as the decomposition in (A). However, if the logic is applied to the computation of the redundant (mutual/shared) part (red) – sum of both mutual information values *I(X; S)* (green and red) and *I(Y; S)* (blue and red) and subtracting the total mutual information I*(X,Y; S)* – the redundant information is not non-negative. Therefore, there must be another part of the diagram that contains the extra information (yellow) that is part of the total information *I(X, Y; S)*.

To derive this measure, we used the Partial Information Decomposition (Williams and Beer, 2010) to decompose the mutual information between a set of variables into redundant, synergistic and unique contributions. The PID distinguished three types of contributions to the information among two variables, or predictors, (here called *A*_*1*_ and *A*_*2*_ for distinction between the formalisms) and a third one, the target *C* (See Schematic in Figure 8A). If *A*_*1*_ and *A*_*2*_ both carry the same information about *C* it suffices to know only one of them to fully determine the mutual information between them and *C*; this type of information is called redundant. If only *A*_*1*_ carries information about *C* it is referred to as unique information; similarly, this holds true for *A*_*2*_. Finally, if neither of the variables alone provide information about *C*, but they do only if observed together, we refer to it as synergistic information. To compute the different terms of the mutual information decomposition, either the synergy or the redundancy suffices, because the remaining terms can be thereafter computed. For example, if we compute the redundancy between *A*_*1*_ and *A*_*2*_, then the unique information carried by each source can be obtained by subtracting the redundancy from the mutual information between the particular source and the target variable *unique*(*A*_*1*_) = *I*(*A*_*1*_; *C*) − *redundancy*(*A*_*1*_, *A*_*2*_). The mutual information *I*(*A*_*1*_; *C*) cannot contain any synergy, because the source *A*_*1*_ is observed independently from *A*_*2*_. Synergy thus becomes the remaining term of the full mutual information *I*(*A*_*1*_, *A*_*2*_; *C*) after subtraction of all the other parts.

**Figure 8.**
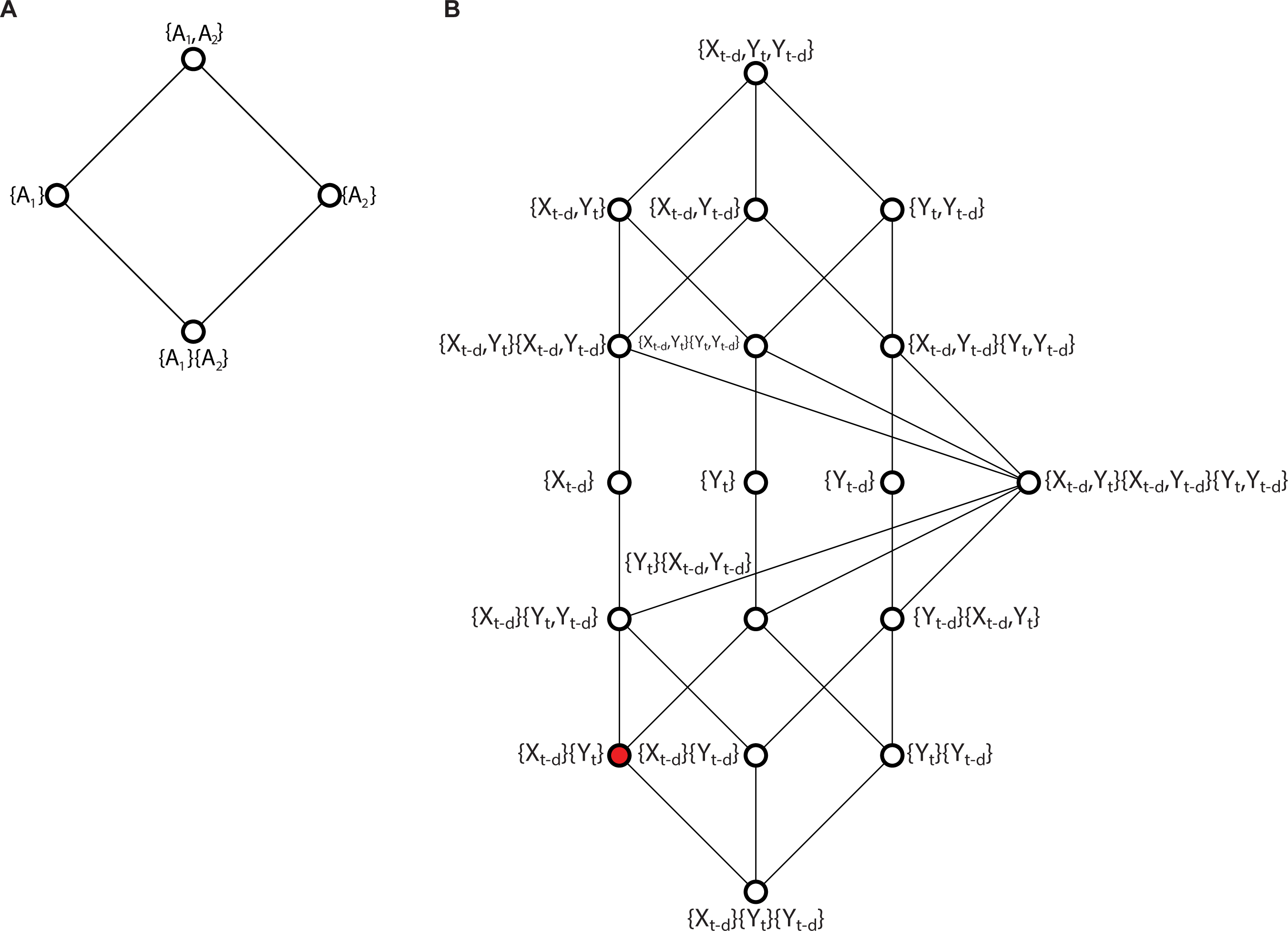
Scheme of the information lattice used to compute the FIT. **A.** Partial information decomposition lattice for 2 sources A_1_ and A_2_. **B.** Partial information decomposition lattice for 3 source (*X*_*t*−*d*_, *Y*_*t*−*d*_, *Y*_*t*_). The lattice is ordered based on the following: a collection of sources A is considered to “succeed” a second collection B (i.e., to be above in the lattice) if for each source in A there exists a source in B that does not provide any additional information than the one in A. The bottom term represents joint redundancy between all three variables and the top their synergy. The FIT measure is marked in red.

Williams and Beer (Williams and Beer, 2010) introduced a measure of redundancy called *I*_*min*_ which quantifies the information associated with a specific outcome *c* provided by a set of possible sources *A* = {*A*_*1*_, *A*_*2*_, …, *A*_*k*_}, and it is defined as:

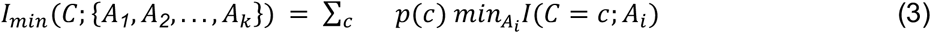

where *I*(*C* = *c*; *A*_*i*_) is a quantity called *specific information*, defined as:

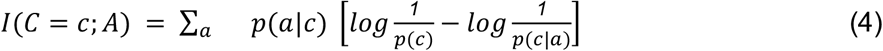

where *a* is one of all possible values of *A*_L_. *I*_*min*_ quantifies redundancy in the form of information that is common to all the sources within *A*: the minimum information that any source provides. The mutual information can be considered a special case of redundancy (i.e., the redundancy of a source with itself). This allows the use of the redundancy measures to compute all terms of the PID. However, the *I*_*min*_ measures themselves do not quantify the unique contribution of a single part of the decomposition. For example, *I*_*min*_(*C*; {*A*_*1*_}), which is equal to the mutual information between *C* and *A*_*1*_, includes the redundant information shared by *A*_*1*_ and *A*_*2*_. Nevertheless, it is possible to reconstruct the information contributed independently by each source of the decomposition from the *I*_*min*_ measures, thanks to the natural ordering among all terms of the decomposition. For example, the redundancy between *A*_*1*_ and *A*_*2*_ would precede the unique contribution of either *A*_*1*_ or *A*_*2*_, because its knowledge is necessary to compute the unique contributions. This ordering creates a lattice from the elements of *A*(*R*) (i.e., the set of all parts of the decomposition based on those sources). In the bivariate case, the terms are defined as follows: the redundant information between sources *A*_*1*_and *A*_*2*_ is written as { *A*_*1*_}{ *A*_*2*_}; the unique information of a source *A*_*1*_ as { *A*_*1*_} and the synergy between *A*_*1*_ and *A*_*2*_ will be noted as { *A*_*1*_, *A*_*2*_}. For the decomposition of the mutual information between two sources *A*_*1*_ and *A*_*2*_ with a target *C* the lowest term in the lattice is the redundancy between the two sources, whereas the terms above include the unique contributions; the highest term incorporates the synergistic effects (Fig. 8A). Within this structure, a higher element in the lattice provides at least as much redundant information as any lower term; the highest element of the lattice then completes all the information that is present in the mutual information between all sources and the target *I*(*R*; *S*). To compute the contribution of each node in the lattice, we need to subtract all information that is provided by underlying nodes. The amount of information provided by a single node is called partial information term ∏_*□*_ and it is defined recursively as (see (Williams and Beer, 2010) for further details):

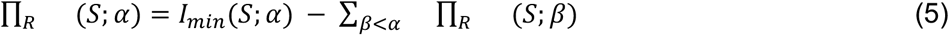

We defined our FIT measure, by applying the above explained PID to the decomposition of an information system made of a set of three predictors *R* = {*X*_*t*−*d*_, *Y*_*t*−*d*_, *Y*_*t*_} and their mutual information with the target stimulus feature *S* (Fig. 8B). The FIT quantifies the information about *S* present in *Y*_*t*_ that is redundant with respect to information already present in *X*_*t*−*d*_ and unique with respect to *Y*_*t*−*d*_. Consequently, the FIT is expressed by the following partial information term (red dot in Fig. 8B):

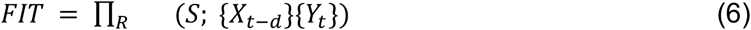

Note that it is not necessary to apply any conditioning on the past of *Y*, because this is implicitly accomplished by excluding the information captured by the lower ∏_*□*_ term of the lattice that includes the past of *Y*, namely ∏_*R*_ (*S*; {*X*_*t*−*d*_}{*Y*_*t*_}{*Y*_*t*−*d*_}.

### Definition and calculation of Directed Feature Information (DFI)

A recent measure called Directed Feature Information (DFI) was introduced to quantify feature-specific directed information (Ince et al., 2015):

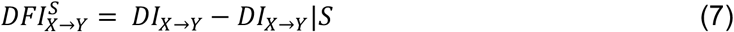

The DFI expresses the difference between the total information transfer between *X* and *Y* (*DI*_*X*→*Y*_) and the information transfer when the stimulus *S* is known (*DI*_*X*→*Y*_|*S*). The first term quantifies the total amount information transferred between *X* and *Y*, whereas the second term quantifies the amount of information that is transferred for a fixed *S*. Hence, the subtraction was considered as the remaining information transfer between *X* and *Y* about *S*. In other words, the DFI quantifies the amount of new information transferred from *X* to *Y* that is related to the variations in *S* (Ince et al., 2015). The DFI can also be expressed as the co-information between *X* and *Y* with respect to *S* (see Supplementary Methods for the full derivation):

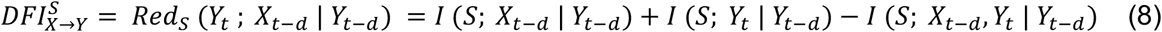

*Red*_*S*_ (*Y*_*t*_; *X*_*t*−*d*_ | *Y*_*t*−*d*_) represents the so-called redundancy (Pola et al., 2003; Schneidman et al., 2003) between *X*_*t*−*d*_and *Y*_*t*_ about stimulus *S* conditional on *Y*_*t*−*d*_. In general, positive redundancy measures the overlap of the MI about the stimulus feature *S* that is present in two different responses. Negative values of redundancy indicate synergy, which occurs when the two variables carry more MI when considered together than alone (i.e., in cases when the stimulus modulates their relationship) (Ince et al., 2015). This occurs whenever the third term is larger than the information carried by the two variables observed independently. Despite the fact that DFI was successfully applied to detect feature-specific transfer (Ince et al., 2015), its property of potentially being negative hinders its interpretation as the amount of information transfer.

To clarify this issue, let us consider all information interactions between two variables *X* and *Y*. The total amount of information in the system is quantified by the joint entropy *H*(*X*, *Y*), whereas the amount of information carried by each variable separately is quantified by the entropy values *H*(*X*) and *X*(*Y*), respectively. The shared part of information is quantified by the mutual information between those variables *I*(*X*; *Y*). In order to quantify information about a single variable *S*, we can replace the entropies with mutual information terms with respect to *S*. Now, the total information about *S* is quantified by the mutual information *I*(*X*, *Y*; *S*) and, consistently, the information separately carried by each variable is given by *I*(*X*; *S*) and *I*(*Y*; *S*), respectively. The shared part of information between the variables is quantified by the redundancy between *X* and *Y* or *Red*_*S*_(*X*; *Y*)(i.e., corresponding to the unconditional version of eq. 7)

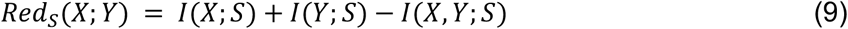

Eq. 9 demonstrated that *Red*_*S*_(*X*; *Y*) can be negative whenever the total amount of information about *S* is larger than the sum of both individual contributions, despite the fact that the shared part of information is included in the sum twice. In such case, the emergent information carried by the interaction between the two variables is called *synergistic* information. Hence, we conclude that the DFI does not satisfy the third predicate, because it does not exclusively represent information that is shared between *X* and *Y*.

### Numerical simulations

We performed numerical simulations to test the reliability of our novel measure, the feature-specific information transfer (FIT). We analyzed different scenarios, each representing particular set of conditions under which feature-specific communication occurs. In each scenario, we simulated a network of two nodes, the first node called *X* (the sender of information) and a second node call *Y* (the receiver of information). Each node generated activity at a given time as a Poisson process with a given time-dependent mean. The time-dependent mean activity had a constant background term simulating ongoing activity intrinsically generated within the node. The representation of stimulus information by the sender node X was modeled with a stimulus-driven term in the time dependent firing rate of X, which was modeled as a Gaussians-shaped bump of activity (with width quantified by the Gaussian’s standard deviation sigma) centered around a given post-stimulus time *t*_*1*_. We assumed that the amplitude of the Gaussian profile was modulated by a stimulus. We assumed we had four different stimuli (the stimulus feature was thus indexed from 1 to 4), and the stimulus to be presented in each trial was drawn at random from a uniform distribution. The baseline activity of both *X* and *Y* was set to 1 spike/sec. Information transfer from *X* to *Y* was achieved by adding the activity of *X* to the baseline signal of *Y* with a communication delay delta.

Simulations were based on three scenarios. The first scenario (Fig. 1A, left panel) simulated a simple stimulus-dependent information transfer from node *X* to *Y*. In this case, node X had one stimulus modulated Gaussian bump of activity which was transferred to Y with a certain delay. Transfer happened only during the stimulus-modulated Gaussian bump. While a transmission of information across areas which is strictly limited to a given window may not be biologically realistic, it was useful in our simulations to check whether the different measures would pick spurious transfer of information in the time windows when there was none. In the second scenario (Fig. 1B, left panel) there was transfer of information that was stimulus-unrelated. To achieve this, the sender node *X* was split into two sub-nodes, called *X*^+^ and *X*^−^. *X*^+^ was simulated as in scenario 1, whereas *X*^−^ was equal to baseline activity minus the Gaussian profile of signal *X*^+^plus a noise drawn from a Poisson distribution with mean equal to the signal value at a given time point. The baseline activity (the activity outside the window with the Gaussian stimulus modulation bump) was set to a constant of 1 spike/sec, and it was identical for both *X*^+^ and *X*^−^. Activities of both sub-nodes were then added together and divided by two (that is, averaged) before being injected into the signal of the receiver *Y*. This process cancels out all stimulus-dependent information transfer to signal *Y*, while leaving stimulus-independent transfer unaffected. The third scenario (Fig. 1C, left panel) was designed to study the effect of an external source of information influencing signal *Y* without passing through the communication channel between *X* to *Y*. To do so, we added information about a second stimulus *S*^*′*^ having the same probability distribution as *S*, but drawn from it independently, to signal *Y*. The Gaussian profile added to the signal *Y* at *t*_*2*_ was thus modulated by the stimulus values *S*^*′*^.

All simulations were run with a 1 ms resolution from −20 to 100 ms peri-stimulus. In figures 1 and 2, and supplementary figure S1 and S2, the onset of the Gaussian stimulus activation bump was set to 20 ms in X and it lasted 20 ms (the standard deviation of the Gaussian bump was 6.6 ms). In the communication window (from 20 to 40 ms), the activity of X was transmitted to Y with a delay of 20 ms. Thus, information arrived in Y starting at 40 ms and Y did not receive more information after 60 ms.

In Figure 3 only, the information transfer window and Gaussian bump (of standard deviation 26.6 ms) lasted 80 ms, starting at the beginning of the bump in the time course in X activity plotted in Fig 3, most leftward panels. In Figure 3, the delay of transmission of this information to Y in the communication window was 5, 15 and 25 ms in panels A, B and C respectively.

In Supplementary Figure 1B, the top panel has simulations taken from the scenario 1 (panel A) similar to Fig 1A, but we added further noise to make it more difficult to detect information transfer. The noise was added by adding a sine wave with a period of 200 ms and random phase at each trial to both X and Y. Since the random phase was independently chosen for X and Y, this addition greatly reduces the influence of X on Y (and thus makes it more difficult to detect information transfer).

In Supplementary Figure 1C, we simulated a case where there is high synergy of stimulus representations between X and Y. This high synergy was obtained by increasing (by a factor of 2.5 with respect to Figure 1) the modulation with the stimulus identification number of the height of the Gaussian bump of stimulus-related activity in X. This larger modulation induces very large stimulus-dependent changes in the way X and Y are correlated (because there is much correlation for stimuli with a higher bump of activity, as X influences more Y for such stimuli than for stimuli that generate only low activity in X). These stimulus dependent correlations generate a large synergy in the joint information about the stimulus carried jointly by X and Y, that is reflected as a large value of the stimulus-dependent component of the mutual information decomposition (that is larger than the other terms), as described by (Pola et al., 2003). We observed that DFI values reported a significant negative peak at the position of the transfer, confirming that DFI values unlike DI and FIT, are decreased by presence of synergy. This is because the measure of redundancy used in the definition of DFI includes actually both redundancy and synergy, and synergy leads to negative DFI values when it is incorrectly subtracted out as redundancy in the definition of DFI (Ince et al., 2015).

For each scenario (Figs 1–3 and Fig S1), we simulated 100 000 trials, except for those used for robustness testing. In Fig S2, to test the robustness of the algorithm for different trial numbers, tests were run on 2^8^ to 2^17^ number of trials.

#### Information theoretical analysis

For every simulation, we computed values of DI, DFI and FIT in a time-resolved manner. To compute the full joint probability distribution required for the calculation of information theoretical measures, we binned all variables at a given time point across trials into three bins, using a technique that distributes equally the number of samples per bin. The number of bins was set to three, as a trade-off between the size of the joint probability distribution and computability cost. Results were however found to be qualitatively similar for other time bin numbers in the region 2 to 10. For all calculations, we computed the FIT, DI and DFI as a function of the parameter time t and delay d. However, for the simulations (in which the real delay used to generate the data was known), we plotted results using the delay value corresponding to the one used to simulate the data. For analysis of real data, we selected for further analyses the range of delays and times for which the transferred information values were significant according to cluster statistics (Maris and Oostenveld, 2007). We refer to the methods for each specific dataset for further details.

To correct for biases due to the limited number of samples, we used the Quadratic Extrapolation procedure (Panzeri et al., 2007; Strong et al., 1998). The bias grows quadratically as a function of the logarithmic decrease in the number of trials. To correct for such bias, we first computed the value of a given information quantity using either all, half or a quarter of the available simulated trials, and then we fitted a quadratic curve. The constant coefficient of the quadratic curve was the corrected value for the information quantity. To increase accuracy in the information theoretic measures, we computed the values for both halves and all quarters and then averaged across them.

#### Statistical analysis

To assess the level of significance of the different information theoretic measures, we used nonparametric permutation techniques. For every time point, we established its 0.1th – 99.9th quantile interval under the null hypothesis. To obtain the values under the null hypothesis, we randomly shuffled stimulus order across trials for DFI and FIT, and values of *X* for DI. In this manner, we computed 100 surrogate values based on independent permutations. To establish the desired percentile values, we fitted a Gaussian distribution to the surrogate data.

### Analysis of Neurophysiological Data

#### EEG and face detection task

##### Experimental conditions and behavioral tasks

In the first set of analyses, we computed the DI, DFI and FIT measures on a publicly available EEG dataset (Rousselet et al., 2014). The data was recorded during a face detection task. Participants (N=16) were presented with an image hidden behind a bubble mask. On half of the trials, the image was a face and the other half contained a random texture. Participants were instructed to determine the content of the image and report whether a face was present or not. For our purposes, we only considered correct trials where the face was correctly detected by the participants (approximately 1000 trials per subject). The dataset contains preprocessed EEG data from the electrodes in the left and the right occipito-temporal regions that display the highest mutual information with the visibility of the contralateral eye. The dataset contains 15 subjects as one was discarded for low quality of the recording.

##### Information theoretic measures

We computed the first derivatives of the EEG signal for both occipito-temporal sensors and used both its absolute values and first derivatives to compute the information quantities. We used both the raw and absolute time derivatives to be consistent with the analyses performed in previous papers (Ince et al., 2016; Rousselet et al., 2014). To establish the joint probability distribution for computation of the information quantities, we binned both the derivatives and absolute values into 3 equally populated bins resulting into 9 possible values for each random variable. The visibility of an eye was considered as the stimulus and it was also binned into 3 bins. We computed the information quantities for all combinations of direction of the transfer (left to right, right to left) and a particular eye (left and right). These values were computed for every subject and then we averaged across them.

##### Statistical analyses

We established significant increase in information measures using a cluster-based nonparametric statistical test introduced in (Maris and Oostenveld, 2007). First, we computed the surrogate values of the information quantities for all time points and delays using 100 permutations of eye visibility. For each of those permutations, we created clusters of values higher than a threshold, based on 8-adjacency in the 2D time-delay grid. We set the threshold to 97.5^th^ quantile, that we obtained by ranking all the values in the grid. Then, we computed cluster level statistic by summing all values of the given information quantity within the cluster. Finally, we took the largest cluster-level statistics for each of the permutations and compared them to cluster-level statistic in the non-shuffled data. We considered as significant those clusters for which cluster-level statistics was lower than the cluster-level statistics of less than 5 clusters obtained from the shuffled data. Hence, we only considered significant those that would score higher than 95th place in a rank test among the cluster-level statistics.

#### MUA and multisensory stimulation

##### Experimental conditions

Multiunit activity (MUA) was recorded from the visual and somatosensory territories of the thalamus and corresponding primary cortical regions (N=6 rats) in three stimulation conditions: visual stimulation, whiskers tactile stimulation and bimodal stimulation (simultaneous visual and tactile, see (Bieler et al., 2018)). The aim of our analyses was to assess the presence of stimulus-specific information transfer from thalamus to cortex. The placement of recording sites enabled simultaneous recording from supragranular (S), granular (G) and infragranular (I) layers of S1 and the ventral posteromedial nucleus (VPM) of the thalamus as well from S, G and I layers of V1 and the dorsal geniculate nucleus (dLGN) of the thalamus. All experiments were conducted during the light phase under sleep-like conditions mimicked by urethane anesthesia (Bitzenhofer et al., 2015). By these means, the interference with spontaneous whisking and the impact of alert state, which modulates cross-modal integration, were avoided. Unimodal (either light flash or whisker deflection) or bimodal (simultaneous light flash and whisker deflection) stimuli were applied using a custom-made stimulation device (Sieben et al., 2013). Briefly, whiskers were stimulated by deflection through compressed air-controlled roundline cylinders gated via solenoid valves. The device produced almost silent (6~10dB), non-electrical stimulation with precise timing (0.013 ± 0.81 ms) that was constant over all trials/conditions. For full eye field visual stimulation, 50-ms-long LED light flashes (300 lux) were used. For bimodal stimulation, whisker deflection and light flashes were applied in the same hemifield. Stimuli were randomly presented across trials in blocks of three different stimulation conditions (unimodal tactile, unimodal visual, bimodal visual-tactile). In our analysis we considered only stimulation contralateral to the recording brain areas. Each type of stimulus was presented 100 times contralateral to the recording electrodes with an interstimulus interval of 6.5 ± 0.5 s. To achieve a physically simultaneous stimulation of whiskers (valve-controlled whisker stimulation) and eyes (instantaneous light flash), the time delay of whisker stimulation was calculated to match visual stimulation onset. The non-stimulated eye was covered with an aluminum foil patch.

The surgery was performed under ketamine/Xylazine anesthesia, the rat’s eyes were covered with ointment (Bepanthen), and the ear canals were filled with silicon adhesive (Kwik-Sil, World Precision Instruments) to block auditory input. Extracellular recordings of the local field potential (LFP) and multiunit activity (MUA) were performed from head-fixed rats under light urethane anesthesia (0.5 g/kg body weight, i.p.; Sigma-Aldrich) using custom made one-shank 1632-channel electrodes (0.5-3 M_Ω Silicon Michigan probes, Neuronexus Technologies; 100-μm intersite spacing) that were angularly inserted into S1 barrel field (ventral-medial, 50°, 2.4-2.6 mm posterior and 5.5-5.8 mm lateral to bregma) and V1 (ventral-rostral, 45°, 6.9-7.1 mm posterior and 3.4-3.7 mm lateral to bregma) to a depth of 5.3 mm and 5.5 mm, respectively. Electrodes were labeled with DiI (1,1’-dioctadecyl-3,3,3’,3’-tetramethylindocarbocyanine; Invitrogen) for postmortem reconstruction of their tracks in histologic sections. A silver wire was inserted into the cerebellum and served as ground and reference electrode. The body temperature of the animal was kept constant at 37°C during recording. The position of recording sites over layers was confirmed by electrophysiological (i.e., reversal of the evoked potential between supragranular and granular layers) and histologic (i.e., granular cell body layer) landmarks. Neural activity was recorded at a sampling rate of 32 kHz using a multichannel extracellular amplifier (no gain, Digital Lynx 10S, Neuralynx) and the acquisition software Cheetah. The signal was bandpass filtered (0.1 Hz and 5 kHz) by the Neuralynx recording system, for antialiasing, and then down-sampled by a factor of 8 obtaining a sampling rate of 8 kHz (Bieler et al., 2018). In the current work, we used the recordings from infragranular layers of S1 and V1 and from VPM and LGN. Data was imported and analyzed offline using custom-written tools in Matlab software (version R2018A (Math-Works), and Multi Unit Activity (MUA) was extracted as in (Safaai et al., 2015). For all channels, spike times were first detected from the band-passed (400–3,000 Hz, fourth-order IIR Butterworth Filter) extracellular potential in each electrode by threshold crossing (>3 SD). A spikes train was obtained for all channels using a temporal binning of 0.125 ms (1/8kHz). For each brain region, mass MUA was obtained by pooling together the trains of all recorded spikes form all electrodes related to that region, a refractory period of 1ms was considered and the resulting train was resampled at 1 KHz (1 ms time bins). Finally, we resized the time bins from 1 ms to 10 ms, summing the number of spikes contained in 10 consecutive time bins of the original signal to obtain a 100 Hz MUA signal.

##### Information theoretic measures and statistical analysis

To establish the joint probability distribution to obtain the information quantities, we binned the values of the MUA signal into 2 equally populated bins resulting into 2 possible values for each random variable. We computed the values of the information quantities (MI, DI, DFI, and FIT) for every subject and then we averaged these values across them. We established significance of the information measures using a cluster-based nonparametric statistical test exactly as we did for the EEG and face detection task. We computed the surrogate values of the information quantities for all time points and delays using 100 permutations of stimuli.

#### Source-level high-gamma MEG activity and visuomotor task

##### Experimental conditions, behavioral tasks and brain data acquisition

The third set of analyses was performed on an MEG dataset collected while participants performed an associative visuomotor mapping task (Brovelli et al., 2017, 2015).The task required participants to perform a finger movement associated to a digit number: digit “1” instructed the execution of the thumb, “2” for the index finger, “3” for the middle finger and so on. Maximal reaction time was 1s. After a fixed delay of 1s following the disappearance of the digit number, an outcome image was presented for 1s and informed the subject whether the response was correct, incorrect, or too late (if the reaction time exceeded 1s). Incorrect and late trials were excluded from the analysis, because they were either absent or very rare (i.e., maximum 2 late trials per session). The next trial started after a variable delay ranging from 2 to 3 s (randomly drawn from a uniform distribution) with the presentation of another visual stimulus. Each participant performed two sessions of 60 trials each (total of 120 trials). Each session included three digits randomly presented in blocks of three trials. The average reaction time was 0.504s ± 0.004s (mean ± s.e.m.). Anatomical T1-weighted MRI images were acquired for all participants using a 3-T whole-body imager equipped with a circular polarized head coil. Magnetoencephalographic (MEG) recordings were performed using a 248 magnetometers system (4D Neuroimaging magnes 3600). Visual stimuli were projected using a video projection and motor responses were acquired using a LUMItouch® optical response keypad with five keys. Presentation® software was used for stimulus delivery and experimental control during MEG acquisition.

##### Brain Atlas: MarsAtlas

Single-subject cortical parcellation was performed using the *MarsAtlas* brain scheme (Auzias et al., 2016). After denoising using a non-local means approach (Coupe et al., 2008), T1-weighted MR-images were segmented using the FreeSurfer “recon-all” pipeline (http://freesurfer.net). Grey and white matter segmentations of each hemisphere were imported into the BrainVisa software and processed using the Morphologist pipeline procedure (http://brainvisa.info). White matter and pial surfaces were reconstructed and triangulated, and all sulci were detected and labeled automatically (Mangin et al., 2004). A parameterization of each hemisphere white matter mesh was performed using the Cortical Surface toolbox (http://www.meca-brain.org/softwares/). It resulted in a 2D orthogonal system defined on the white matter mesh, constrained by a set of primary and secondary sulci (Auzias et al., 2016).

##### Single-trial high-gamma activity (HGA) in MarsAtlas

MEG signals were down-sampled to 1 kHz, low-pass filtered to 250 Hz and segmented into epochs aligned on finger movement (i.e., button press). Epoch segmentation was also performed on stimulus onset and the data from −0.5 and −0.1 s prior to stimulus presentation was taken as baseline activity for the calculation of the single-trial high-gamma activity (HGA). Artefact rejection was performed semi-automatically and by visual inspection. For each movement-aligned epoch and channel, the signal variance and z-value were computed over time and taken as relevant metrics for the identification of artefact epochs. All trials with a variance greater than 1.5*10-24 across channels were excluded from further search of artefacts. Metrics such as the z-score, absolute z-score, and range between the minimum and maximum values were also inspected to detect artefacts. Channels and trials displaying outliers were removed. Two MEG sensors were excluded from the analysis for all subjects.

Spectral density estimation was performed using multi-taper method based on discrete prolate spheroidal (slepian) sequences (Swift et al., 1995). To extract high-gamma activity from 60 to 120Hz, MEG time series were multiplied by *k* orthogonal tapers (*k* = 8) (0.15s in duration and 60Hz of frequency resolution, each stepped every 0.005s), centered at 90Hz and Fourier-transformed. Complex-valued estimates of spectral measures 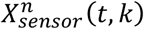, including cross-spectral density matrices, were computed at the sensor level for each trial *n*, time *t* and taper *k*. Source analysis requires a physical forward model or leadfield, which describes the electromagnetic relation between sources and MEG sensors. The leadfield combines the geometrical relation of sources (dipoles) and sensors with a model of the conductive medium (i.e., the head model). For each participant, we generated a head model using a single-shell model constructed from the segmentation of the cortical tissue obtained from individual MRI scans (Nolte, 2003). Leadfields were not normalized. Sources were placed in the single-subject volumetric parcellation regions. For each region, we computed the number of sources *nSP* as the ratio of the volume and the volume of a sphere of radius equal to 3 mm. The k-means algorithm (Tou and Gonzalez, 1974) was then used to partition the 3D coordinates of the voxels within a given volumetric region into *nS* clusters. The sources were placed at the center of each partition for each brain region. The head model, source locations and the information about MEG sensor position for both models were combined to derive single-participant leadfields. The orientation of cortical sources was set perpendicular to the cortical surface, whereas the orientation for subcortical sources was left unconstrained.

We used adaptive linear spatial filtering (Van Veen et al., 1997) to estimate the power at the source level. In particular, we employed the Dynamical Imaging of Coherent Sources (DICS) method, a beamforming algorithm for the tomographic mapping in the frequency domain (Gross et al., 2001), which is a well suited for the study of neural oscillatory responses based on single-trial source estimates of band-limited MEG signals (Hansen et al., 2015). At each source location, DICS employs a spatial filter that passes activity from this location with unit gain while maximally suppressing any other activity. The spatial filters were computed on all trials for each time point and session, and then applied to single-trial MEG data. DICS allows the estimate of complex-value spectral measures at the source level, 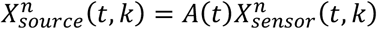, where *A*(*t*) is the spatial filter that transforms the data from the sensor to source level and 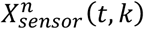 is the complex-valued estimates of spectral measures, including cross-spectral density matrices, computed at the sensor level for each trial *n*, time *t* and taper *k* (for a detailed description of a similar approach see (Hipp et al., 2011)). The single-trial high-gamma power at each source location was estimated by multiplying the complex spectral estimates with their complex conjugate, and averaged over tapers *k*, 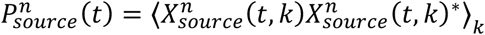, where angle brackets refer to the average across tapers and * to the complex conjugate. Single-trial power estimates aligned with movement and stimulus onset were log-transformed to make the data approximate Gaussian and low-pass filtered at 50Hz to reduce noise. Single-trial mean power and standard deviation in a time window from −0.5 and −0.1 s prior to stimulus onset was computed for each source and trial, and used to z-transform single-trial movement-locked power time courses. Similarly, single-trial movement-locked power time courses were log-transformed and z-scored with respect to baseline period, to produce HGAs for the prestimulus period from −1.6 to - 0.1 s with respect to stimulation for subsequent functional connectivity analysis. Finally, single-trial HGA for each brain region of *MarsAtlas* was computed as the mean z-transformed power values averaged across all sources within the same region.

##### Information theoretic measures

The goal of the MEG analysis was to assess the dynamics of visuomotor-related modulations in DI, DFI and FIT as they evolve over time. The DFI and FIT were thus computed by taking as feature the visuomotor-related *versus* baseline HGA. The visuomotor-related activity was considered as the HGA aligned with movement onset (from −1s to 0.75s), whereas the baseline activity was defined as the HGA at −0.5s prior to stimulus onset. To estimate the joint probability distribution at each time stamp around movement onset, we concatenated the HGA in windows of 20 ms (4 time instants), trials and subjects, and we binned the values into three equally-populated bins, thus resulting into 6 possible values for each random variable. The DI, DF and FIT measures were thus computed in a time-resolved manner on all time points aligned on motor response. We computed information theoretic measures to quantify the information transfer about visuomotor processing between three regions of interest: the medial superior parietal cortex (SPCm), the dorsolateral premotor cortex (PMdl) and the dorsal Inferior Parietal Cortex (IPCd).

## Author contribution

SP and JB conceived the FIT. SP and AB jointly supervised the study. JB implemented the FIT and performed simulations and EEG analysis. VDF performed MUA analysis. AB performed MEG analysis. MB and IO recorded MUA data. AB recorded MEG data. DC contributed methods. SP, AB, JB wrote the paper with contributions from DC and VDF. All authors commented on the manuscript.

## Acknowledgements

This research was supported by the NIH Brain Initiative (grants U19NS107464 and NS108410 to SP), the Simons Foundation (SFARI Explorer 602849 to SP), by the European Research Council (ERC-2015-CoG 681577 to I.L.H.-O.) and the German Research Foundation (Ha 4466/10-1, SPP 1665, SFB 936 B5 to I.L.H.-O). AB was supported by the French National Agency (ANR-18-CE28-0016-01) and the FLAG-ERA (ANR-17-HBPR-0001-02). We thank G. Bondanelli and A. Tlaie Boria for useful comments on an earlier version of this manuscript.

## Supplementary Methods

### Derivation of the DFI as redundancy

Here we report briefly the equations expressing the DFI as a redundancy between mutual information, see (Ince et al., 2015) for more details.

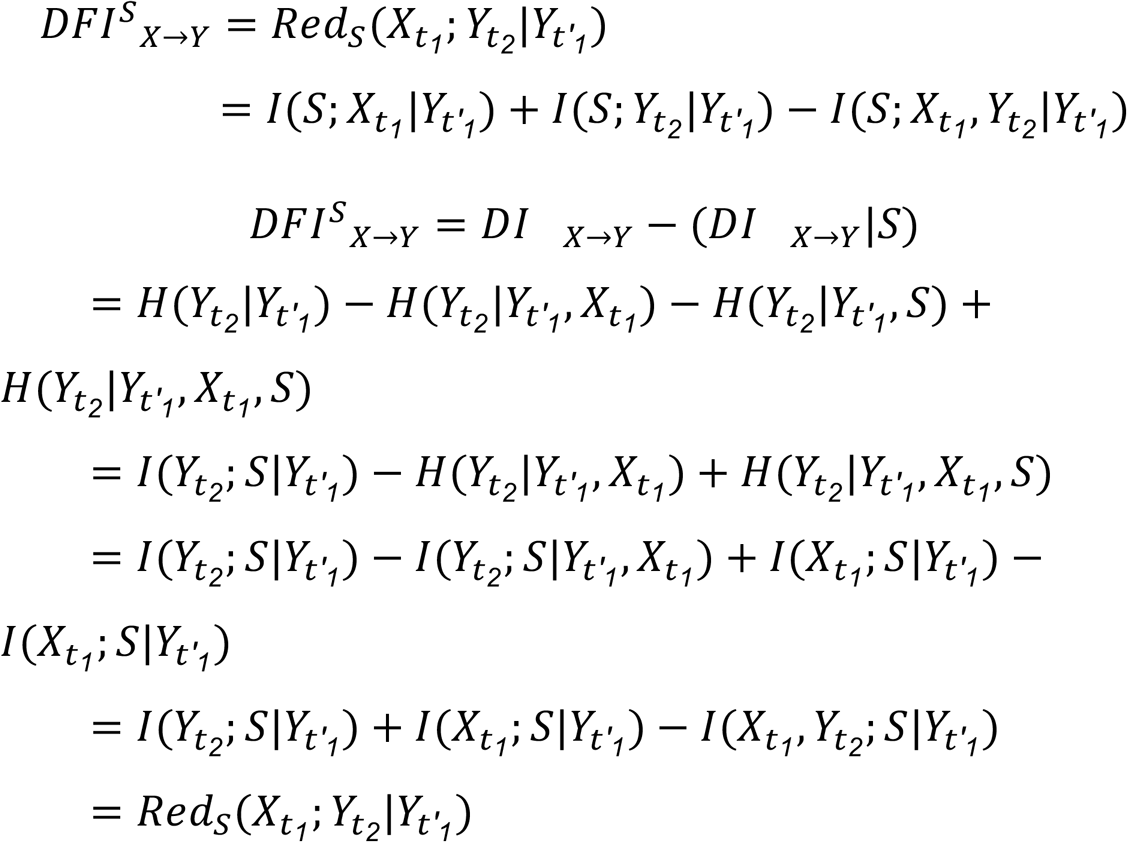

### DFI decomposition according to the PID terms

In order to investigate the relationship between DFI and FIT, we reformulate the DFI as a sum of the partial information terms from the PID, developed by Williams and Beer (Williams and Beer, 2010).

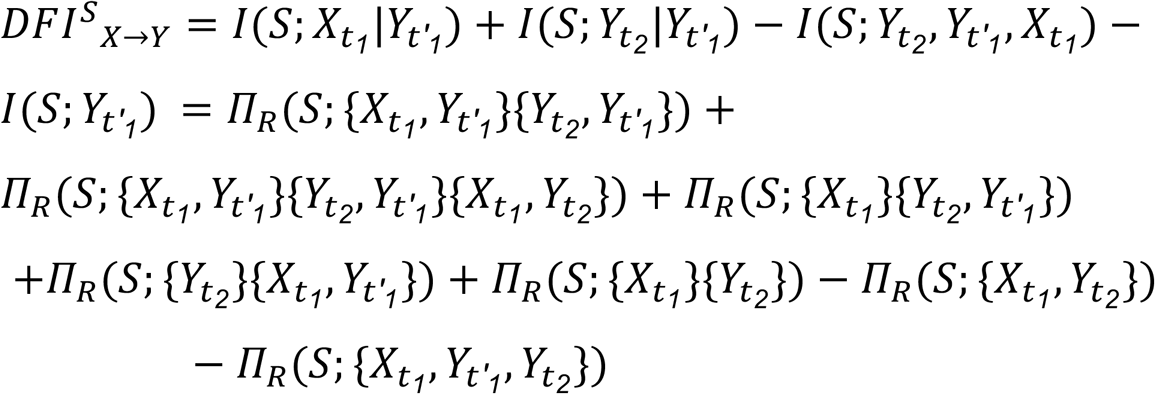

Note that all terms representing a form of redundant information are positive and those representing synergistic information are negative. Therefore, this decomposition demonstrates that the DFI is redundant information minus synergistic information. However, the decomposition does not include only terms regarding the relationship between the past of X and present of Y, but also more complex terms whose intuitive explanation is not trivial. In conclusion, such decomposition shows in which cases the DFI provides appropriate measures of stimulus-specific information transfer.

---

**Supplementary Figure 1.**
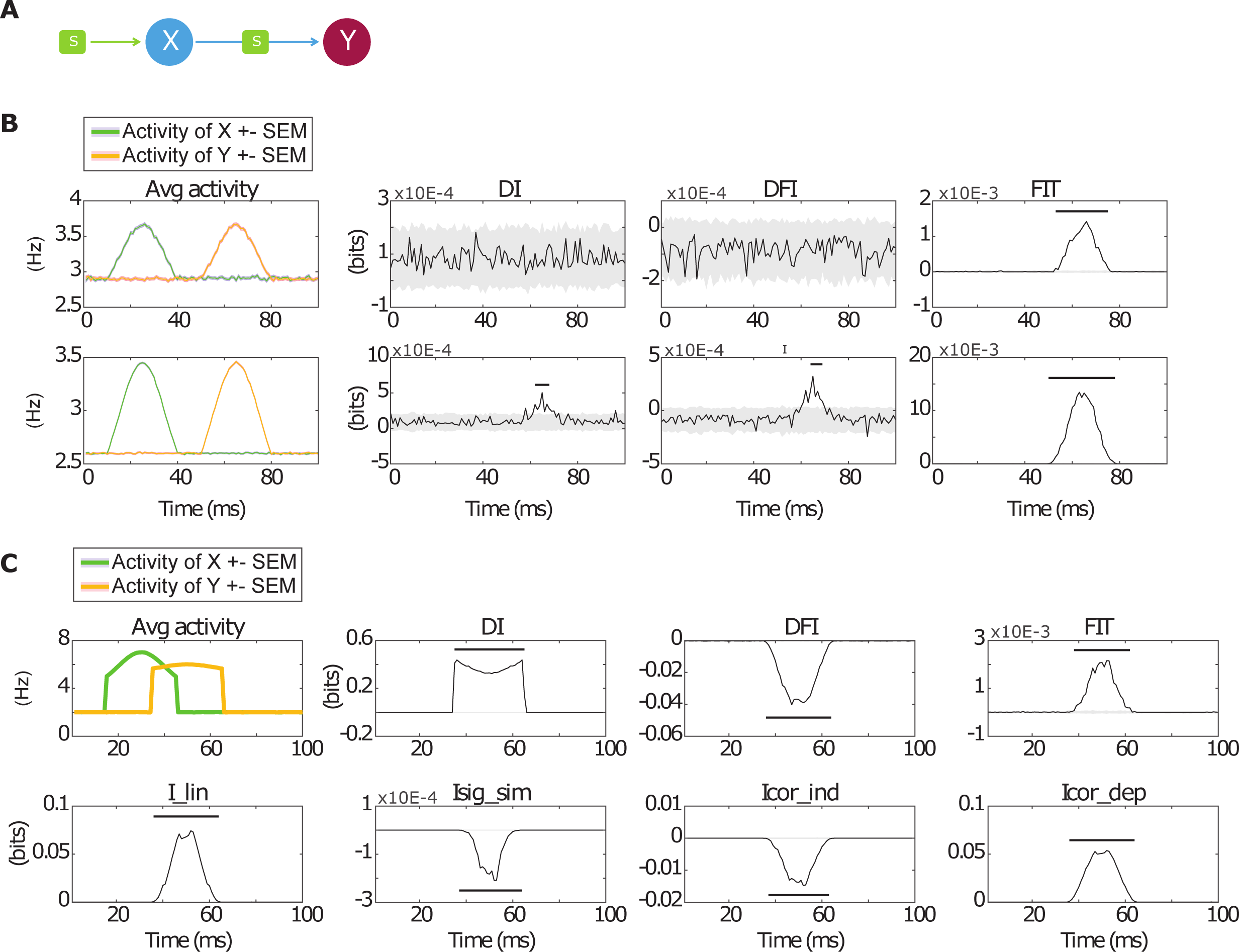
Test of information flow measures with varying degrees of noise and of synergy between nodes. **A.** Scheme of the simulated feed-forward feature-specific communication from X to Y used to test robustness of the measures against noise. **B**. To test noise dependence, we introduced an extra noise into the signal. The results clearly show that DI and DFI are more susceptible to noise as their value was not significant over the whole simulation period in case of high noise (top) and creating a significant peak with much smaller width than the transfer in case of low noise (bottom). FIT (right panels) was able to correctly detect the transfer in both cases, suggesting that it is more robust to noise than DI (left panels) and DFI (central panels). **C.** To confirm the hypothesis the DFI becomes negative due to synergistic effects, we developed a simulation based on the basic transfer scenario **A** by setting the parameters of the time course in such a way to introduce synergy between X and y. This is confirmed by presence of negative signal similarity and positive stimulus dependent correlations (that dominate the stimulus independent correlations) as described by (Pola et al., 2003). It can be observed that DFI values (central panels) created a significant negative peak at the position of the transfer, confirming that DFI values unlike DI (left panels) and FIT (right panels), are decreased by presence of synergy.

**Supplementary Figure 2.**
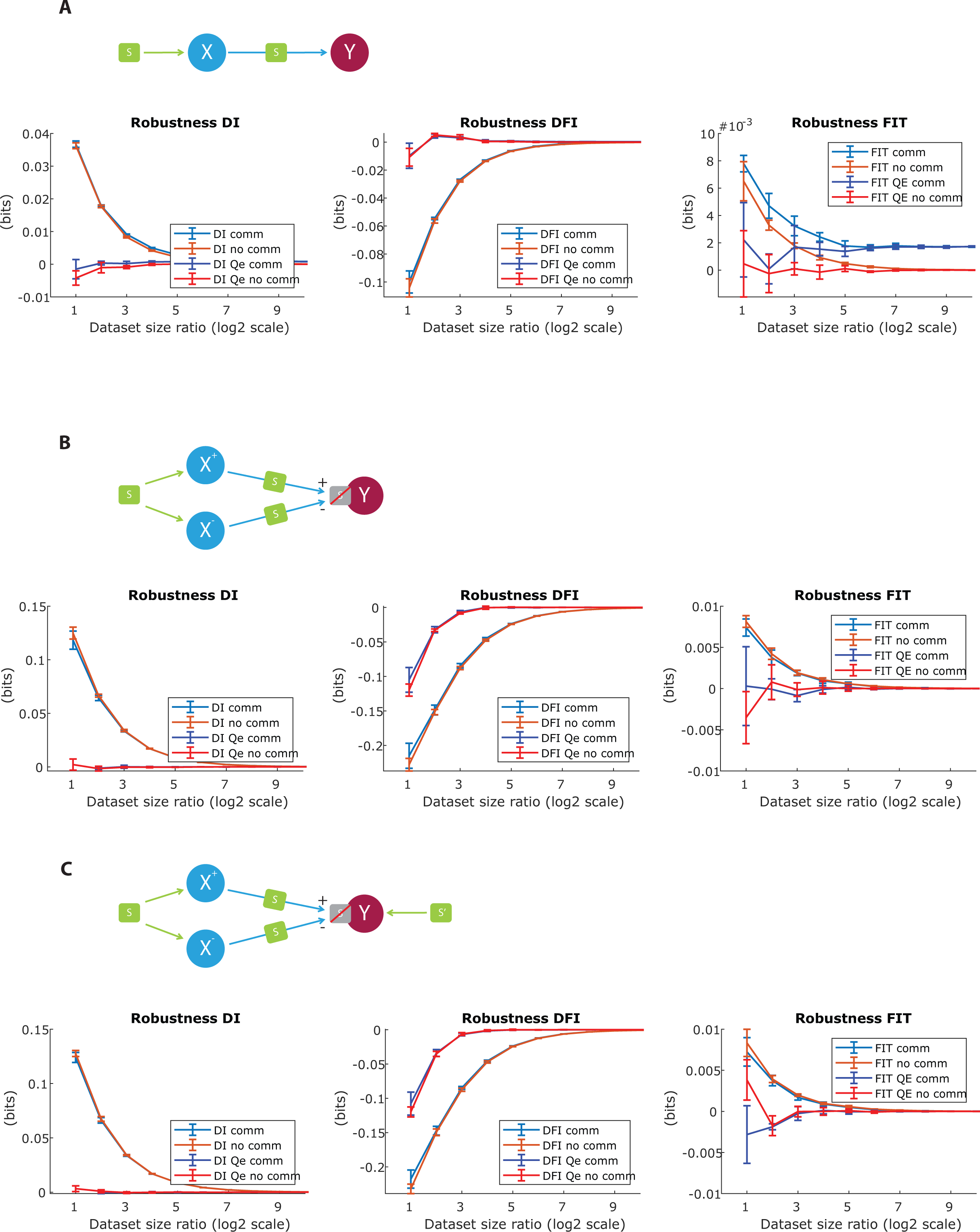
Robustness of FIT and DFI with respect to the finite sampling bias, which is a well-documented problem for information theoretical quantities (Panzeri et al., 2007; Panzeri and Treves, 1996). We tested the measures in all three connectivity scenarios (A, B and C panels) as described in Fig. 1, for time points where transfer did (comm) and did not (no comm) occur and compared an uncorrected and corrected computation of them. For correction, we used the Quadratic Extrapolation (QE) correction (Panzeri et al., 2007; Strong et al., 1998). It can be observed that all quantities behave as generally expected by information theoretic quantities, that is they show there is a bias that grows quadratically with decrease in the logarithm of the dataset size. Moreover, the QE correction was able to decrease the bias considerably.

**Supplementary Figure 3.**
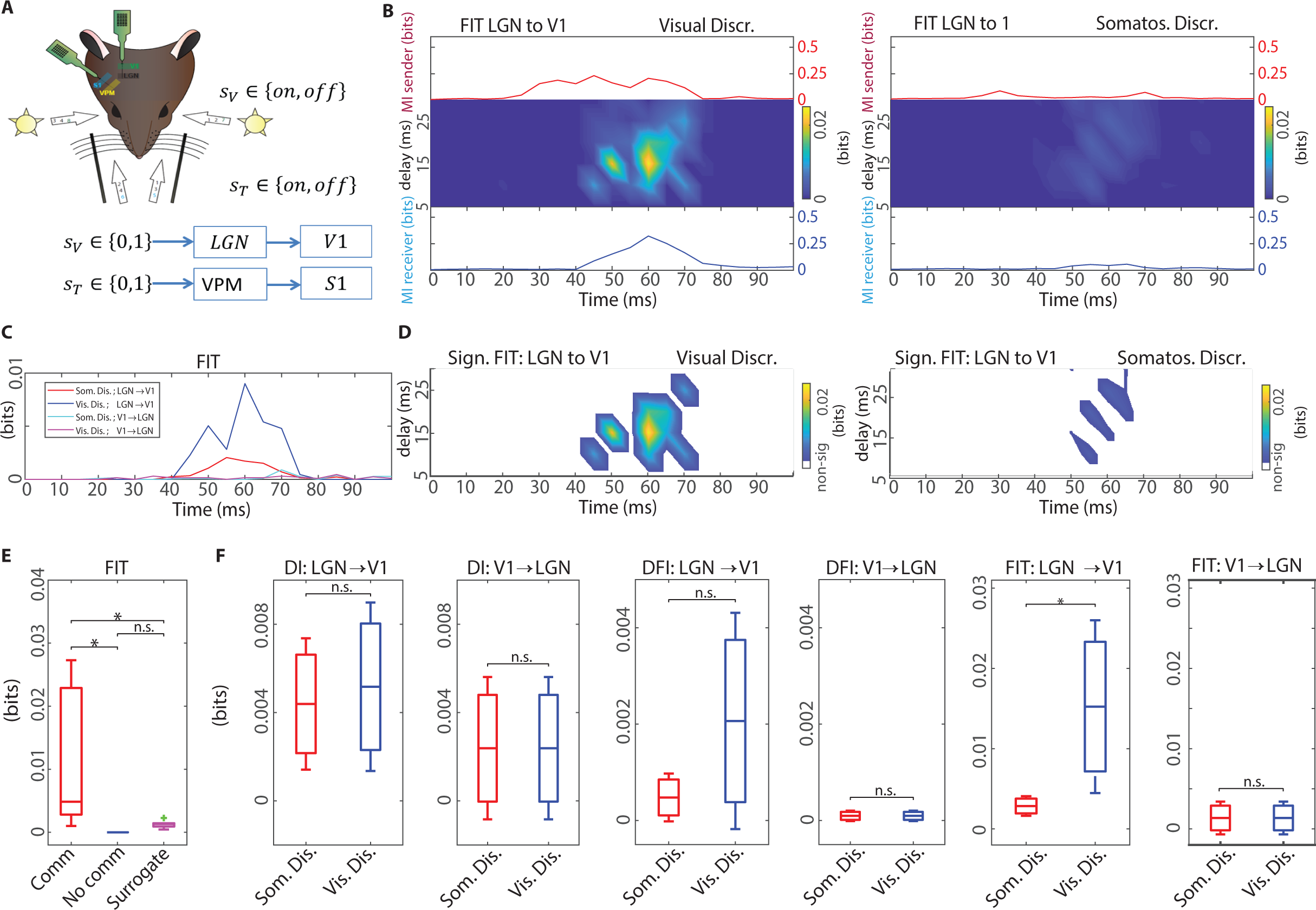
Sensory-related information transfer in the visual system (rat MUA). **A.** Schematic of the experiment. MUA was recorded from the visual and somatosensory territories of the thalamus and associated primary cortical regions in rats. We considered only sensory stimulation contralateral to the recorded regions. **B.** FIT from LGN to V1. The image plots shows the values of FIT (mean across subjects) for each value of delay and post-stimulus time. Line plots above and below show Mutual Information (MI) between the presented stimulus and the recorded MUA in the LGN and V1, respectively. The left panel reports values of information and FIT about the visual discriminative stimulus set, whereas the right panel reports values of information and FIT about the somatosensory-discriminative stimulus set. Information flow from visual thalamus to visual cortex is only about the visual discriminative feature. **C.** FIT values averaged across delays for all combinations of transfer direction and stimulation type. **D.** Image plot (as function of the parameters communication delay and post-stimulus time) of the values of the FIT that are significant (p < 0.05) after a cluster-based nonparametric permutation test (see Methods). The (time,delay) region with significant values of FIT was defined to be the communication window. The left panel reports FIT about the visual discriminative stimulus set, whereas the right panels report FIT about the somatosensory-discriminative stimulus set. **E.** Comparison of FIT values averaged within the (time, delay) communication cluster (indicated as “comm”) with the values obtained averaging FIT within a cluster of the same shape and size, but positioned elsewhere in a prestimulus time window and long delay region in which communication was neither significant nor possible (“no comm”), and surrogate FIT values obtained by random shuffling of the stimulus values**. F.** Comparisons between all 4 combinations of transfer direction (from thalamus to cortex and vice-versa) and conditions (Somatosensory and visually discriminative), for all the three measures. In Panels B-D, we plot and compute the statistical significance of the average across subjects of the information values. In panel E boxplots represent distributions of information values over subjects.

## References

Amblard P-O, Michel OJJ. 2011. On directed information theory and Granger causality graphs. J Comput Neurosci 30:7–16. doi:10.1007/s10827-010-0231-x

Auzias G, Coulon O, Brovelli A. 2016. MarsAtlas: A cortical parcellation atlas for functional mapping. Hum Brain Mapp 37:1573–1592. doi:10.1002/hbm.23121

Barrett AB. 2015. Exploration of synergistic and redundant information sharing in static and dynamical Gaussian systems. Phys Rev E - Stat Nonlinear, Soft Matter Phys 91. doi:10.1103/PhysRevE.91.052802

Bentin S, Allison T, Puce A, Perez E, McCarthy G. 1996. Electrophysiological studies of face perception in humans. J Cogn Neurosci 8:551–565. doi:10.1162/jocn.1996.8.6.551

Bertschinger N, Rauh J, Olbrich E, Jost J, Ay N. 2014. Quantifying unique information. Entropy 16:2161–2183. doi:10.3390/e16042161

Besserve M, Lowe SC, Logothetis NK, Schölkopf B, Panzeri S. 2015. Shifts of Gamma Phase across Primary Visual Cortical Sites Reflect Dynamic Stimulus-Modulated Information Transfer. PLoS Biol 13:1–29. doi:10.1371/journal.pbio.1002257

Bieler M, Xu X, Marquardt A, Hanganu-Opatz IL. 2018. Multisensory integration in rodent tactile but not visual thalamus. Sci Rep 8:1–18. doi:10.1038/s41598-018-33815-y

Bitzenhofer SH, Sieben K, Siebert KD, Spehr M, Hanganu-Opatz IL. 2015. Oscillatory activity in developing prefrontal networks results from theta-gamma-modulated synaptic inputs. Cell Rep 11:486–497. doi:10.1016/j.celrep.2015.03.031

Bosman CA, Schoffelen JM, Brunet N, Oostenveld R, Bastos AM, Womelsdorf T, Rubehn B, Stieglitz T, De Weerd P, Fries P. 2012. Attentional Stimulus Selection through Selective Synchronization between Monkey Visual Areas. Neuron 75:875–888. doi:10.1016/j.neuron.2012.06.037

Bressler SL, Menon V. 2010. Large-scale brain networks in cognition: emerging methods and principles. Trends Cogn Sci 14:277–290. doi:10.1016/j.tics.2010.04.004

Bressler SL, Seth AK. 2011. Wiener-Granger Causality: A well established methodology. Neuroimage. doi:10.1016/j.neuroimage.2010.02.059

Brovelli A, Badier JM, Bonini F, Bartolomei F, Coulon O, Auzias G. 2017. Dynamic reconfiguration of visuomotor-related functional connectivity networks. J Neurosci 37:839–853. doi:10.1523/JNEUROSCI.1672-16.2016

Brovelli A, Chicharro D, Badier JM, Wang H, Jirsa V. 2015. Characterization of cortical networks and corticocortical functional connectivity mediating arbitrary visuomotor mapping. J Neurosci 35:12643–12658. doi:10.1523/JNEUROSCI.4892-14.2015

Brovelli A, Ding M, Ledberg A, Chen Y, Nakamura R, Bressler SL. 2004. Beta oscillations in a large-scale sensorimotor cortical network: Directional influences revealed by Granger causality. Proc Natl Acad Sci U S A 101:9849–9854. doi:10.1073/pnas.0308538101

Cheyne D, Ferrari P. 2013. MEG studies of motor cortex gamma oscillations: Evidence for a gamma “fingerprint” in the brain? Front Hum Neurosci. doi:10.3389/fnhum.2013.00575

Coupe P, Yger P, Prima S, Hellier P, Kervrann C, Barillot C. 2008. An optimized blockwise nonlocal means denoising filter for 3-D magnetic resonance images. IEEE Trans Med Imaging 27:425–441. doi:10.1109/TMI.2007.906087

Crone NE, Sinai A, Korzeniewska A. 2006. Chapter 19 High-frequency gamma oscillations and human brain mapping with electrocorticography. Prog Brain Res. doi:10.1016/S0079-6123(06)59019-3

Darvas F, Scherer R, Ojemann JG, Rao RP, Miller KJ, Sorensen LB. 2010. High gamma mapping using EEG. Neuroimage 49:930–938. doi:10.1016/j.neuroimage.2009.08.041

Emiliani V, Cohen AE, Deisseroth K, Häusser M. 2015. All-optical interrogation of neural circuits. J Neurosci. doi:10.1523/JNEUROSCI.2916-15.2015

Granger CWJ. 1969. Investigating Causal Relations by Econometric Models and Cross-spectral Methods. Econometrica 37:424. doi:10.2307/1912791

Griffith V, Koch C. 2012. Quantifying synergistic mutual information.

Gross J, Kujala J, Hämäläinen M, Timmermann L, Schnitzler A, Salmelin R. 2001. Dynamic imaging of coherent sources: Studying neural interactions in the human brain. Proc Natl Acad Sci U S A 98:694–699. doi:10.1073/pnas.98.2.694

Hansen ECA, Battaglia D, Spiegler A, Deco G, Jirsa VK. 2015. Functional connectivity dynamics: Modeling the switching behavior of the resting state. Neuroimage 105:525–535. doi:10.1016/j.neuroimage.2014.11.001

Harder M, Salge C, Polani D. 2013. Bivariate measure of redundant information. Phys Rev E - Stat Nonlinear, Soft Matter Phys 87. doi:10.1103/PhysRevE.87.012130

Hipp JF, Engel AK, Siegel M. 2011. Oscillatory synchronization in large-scale cortical networks predicts perception. Neuron 69:387–396. doi:10.1016/j.neuron.2010.12.027

Ince R. 2017. Measuring Multivariate Redundant Information with Pointwise Common Change in Surprisal. Entropy 19:318. doi:10.3390/e19070318

Ince RAA, Jaworska K, Gross J, Panzeri S, Van Rijsbergen NJ, Rousselet GA, Schyns PG. 2016. The Deceptively Simple N170 Reflects Network Information Processing Mechanisms Involving Visual Feature Coding and Transfer Across Hemispheres. Cereb Cortex 26:4123–4135. doi:10.1093/cercor/bhw196

Ince RAA, Van Rijsbergen NJ, Thut G, Rousselet GA, Gross J, Panzeri S, Schyns PG. 2015. Tracing the Flow of Perceptual Features in an Algorithmic Brain Network. Sci Rep 5:1–17. doi:10.1038/srep17681

Jerbi K, Ossandón T, Hamamé CM, Senova S, Dalal SS, Jung J, Minotti L, Bertrand O, Berthoz A, Kahane P, Lachaux JP. 2009. Task-related gamma-band dynamics from an intracerebral perspective: Review and implications for surface EEG and MEG. Hum Brain Mapp. doi:10.1002/hbm.20750

Lachaux JP, Axmacher N, Mormann F, Halgren E, Crone NE. 2012. High-frequency neural activity and human cognition: Past, present and possible future of intracranial EEG research. Prog Neurobiol. doi:10.1016/j.pneurobio.2012.06.008

Li M, Han Y, Aburn MJ, Breakspear M, Poldrack RA, Shine JM, Lizier JT. 2019. Transitions in information processing dynamics at the whole-brain network level are driven by alterations in neural gain. PLoS Comput Biol 15:e1006957. doi:10.1371/journal.pcbi.1006957

Mangin JF, Rivière D, Cachia A, Duchesnay E, Cointepas Y, Papadopoulos-Orfanos D, Scifo P, Ochiai T, Brunelle F, Régis J. 2004. A framework to study the cortical folding patternsNeuroImage. doi:10.1016/j.neuroimage.2004.07.019

Maris E, Oostenveld R. 2007. Nonparametric statistical testing of EEG- and MEG-data. J Neurosci Methods 164:177–190. doi:10.1016/j.jneumeth.2007.03.024

Massey JL. 1990. CAUSALITY, FEEDBACK AND DIRECTED INFORMATION James L. Massey. Technology.

Nigam S, Pojoga S, Dragoi V. 2019. Synergistic Coding of Visual Information in Columnar Networks. Neuron 104:402–411.e4. doi:10.1016/j.neuron.2019.07.006

Nolte G. 2003. The magnetic lead field theorem in the quasi-static approximation and its use for magnetoenchephalography forward calculation in realistic volume conductors. Phys Med Biol 48:3637–3652. doi:10.1088/0031-9155/48/22/002

Panzeri S, Harvey CD, Piasini E, Latham PE, Fellin T. 2017. Cracking the Neural Code for Sensory Perception by Combining Statistics, Intervention, and Behavior. Neuron 93:491–507. doi:10.1016/j.neuron.2016.12.036

Panzeri S, Senatore R, Montemurro MA, Petersen RS. 2007. Correcting for the sampling bias problem in spike train information measures. J Neurophysiol 98:1064–1072. doi:10.1152/jn.00559.2007

Panzeri S, Treves A. 1996. Analytical estimates of limited sampling biases in different information measures. Netw Comput Neural Syst 7:87–107. doi:10.1088/0954-898X/7/1/006

Pica G, Piasini E, Safaai H, Runyan C, Harvey C, Diamond M, Kayser C, Fellin T, Panzeri S. 2017. Quantifying how much sensory information in a neural code is relevant for behavior In: Guyon I, Luxburg U V, Bengio S, Wallach H, Fergus R, Vishwanathan S, Garnett R, editors. Advances in Neural Information Processing Systems 30. Curran Associates, Inc. pp. 3686–3696.

Pica G, Soltanipour M, Panzeri S. 2019. Using intersection information to map stimulus information transfer within neural networks. BioSystems 185:104028. doi:10.1016/j.biosystems.2019.104028

Pola G, Thiele A, Hoffmann KP, Panzeri S. 2003. An exact method to quantify the information transmitted by different mechanisms of correlational coding. Netw Comput Neural Syst 14:35–60. doi:10.1088/0954-898X/14/1/303

Ray S, Maunsell JHR. 2011. Different origins of gamma rhythm and high-gamma activity in macaque visual cortex. PLoS Biol 9. doi:10.1371/journal.pbio.1000610

Rousselet GA, Ince RAA, van Rijsbergen NJ, Schyns PG. 2014. Eye coding mechanisms in early human face event-related potentials. J Vis 14. doi:10.1167/14.13.7

Runyan CA, Piasini E, Panzeri S, Harvey CD. 2017. Distinct timescales of population coding across cortex. doi:10.1038/nature23020

Safaai H, Neves R, Eschenko O, Logothetis NK, Panzeri S. 2015. Modeling the effect of locus coeruleus firing on cortical state dynamics and single-trial sensory processing. Proc Natl Acad Sci U S A 112:12834–12839. doi:10.1073/pnas.1516539112

Safaai H, Onken A, Harvey CD, Panzeri S. 2018. Information estimation using nonparametric copulas. Phys Rev E 98:53302. doi:10.1103/PhysRevE.98.053302

Schneidman E, Bialek W, Berry MJ. 2003. Synergy, Redundancy, and Independence in Population Codes. J Neurosci 23:11539–11553. doi:10.1523/jneurosci.23-37-11539.2003

Schreiber T. 2000. Measuring information transfer. Phys Rev Lett 85:461–464. doi:10.1103/PhysRevLett.85.461

Seth AK, Barrett AB, Barnett L. 2015. Granger causality analysis in neuroscience and neuroimaging. J Neurosci 35:3293–3297. doi:10.1523/JNEUROSCI.4399-14.2015

Sheikhattar A, Miran S, Liu J, Fritz JB, Shamma SA, Kanold PO, Babadi B. 2018. Extracting neuronal functional network dynamics via adaptive Granger causality analysis. Proc Natl Acad Sci U S A 115:E3869–E3878. doi:10.1073/pnas.1718154115

Sieben K, Röder B, Hanganu-Opatz IL. 2013. Oscillatory entrainment of primary somatosensory cortex encodes visual control of tactile processing. J Neurosci 33:5736–5749. doi:10.1523/JNEUROSCI.4432-12.2013

Stramaglia S, Angelini L, Wu G, Cortes JM, Faes L, Marinazzo D. 2016. Synergetic and Redundant Information Flow Detected by Unnormalized Granger Causality: Application to Resting State fMRI. IEEE Trans Biomed Eng 63:2518–2524. doi:10.1109/TBME.2016.2559578

Strong SP, Koberle R, De Ruyter Van Steveninck RR, Bialek W. 1998. Entropy and information in neural spike trains. Phys Rev Lett 80:197–200. doi:10.1103/PhysRevLett.80.197

Swift AL, Percival DA, Walden AT. 1995. Spectral Analysis for Physical Applications: Multipaper and Conventional Techniques. J R Stat Soc Ser A (Statistics Soc 158:198. doi:10.2307/2983429

Tou JT, Gonzalez RC. 1974. Pattern recognition principles. Addison-Wesley Pub. Co.

Van Kerkoerle T, Self MW, Dagnino B, Gariel-Mathis MA, Poort J, Van Der Togt C, Roelfsema PR. 2014. Alpha and gamma oscillations characterize feedback and feedforward processing in monkey visual cortex. Proc Natl Acad Sci U S A 111:14332–14341. doi:10.1073/pnas.1402773111

Van Veen BD, Van Drongelen W, Yuchtman M, Suzuki A. 1997. Localization of brain electrical activity via linearly constrained minimum variance spatial filtering. IEEE Trans Biomed Eng 44:867–880. doi:10.1109/10.623056

Van Vugt B, Dagnino B, Vartak D, Safaai H, Panzeri S, Dehaene S, Roelfsema PR. 2018. The threshold for conscious report: Signal loss and response bias in visual and frontal cortex. Science (80-) 360:537–542. doi:10.1126/science.aar7186

Varela F, Lachaux JP, Rodriguez E, Martinerie J. 2001. The brainweb: Phase synchronization and large-scale integration. Nat Rev Neurosci 2:229–239. doi:10.1038/35067550

Vicente R, Wibral M, Lindner M, Pipa G. 2011. Transfer entropy-a model-free measure of effective connectivity for the neurosciences. J Comput Neurosci 30:45–67. doi:10.1007/s10827-010-0262-3

Vinck M, Huurdeman L, Bosman CA, Fries P, Battaglia FP, Pennartz CMA, Tiesinga PH. 2015. How to detect the Granger-causal flow direction in the presence of additive noise? Neuroimage 108:301–318. doi:10.1016/j.neuroimage.2014.12.017

Wibral M, Priesemann V, Kay JW, Lizier JT, Phillips WA. 2017. Partial information decomposition as a unified approach to the specification of neural goal functions. Brain Cogn 112:25–38. doi:10.1016/j.bandc.2015.09.004

Wiener N. 1956. The Theory of Prediction. Mod Math Eng 58:323–329.

Williams PL, Beer RD. 2010. Nonnegative Decomposition of Multivariate Information.

